# Rapid Prediction of Lipid Interaction Sites on Pleckstrin Homology Domains Using Deep Graph Neural Networks and Molecular Dynamics Simulations

**DOI:** 10.1101/2023.12.22.573003

**Authors:** Kyle I.P. Le Huray, Frank Sobott, He Wang, Antreas C. Kalli

**Affiliations:** School of Molecular and Cellular Biology, Faculty of Biological Sciences, University of Leeds, Leeds, UK; Astbury Centre for Structural and Molecular Biology, Faculty of Biological Sciences, University of Leeds, Leeds, UK; Leeds Institute of Cardiovascular and Metabolic Medicine, School of Medicine, University of Leeds, Leeds, UK; Department of Computer Science, University College London, London, UK

## Abstract

Interactions between membrane proteins and specific lipid molecules play a major role in cellular biology, but characterizing these interactions can be challenging due to the complexity and physicochemical properties of membranes. Molecular dynamics (MD) simulations allow researchers to predict protein-lipid interaction sites and generate testable models. MD simulations are however computationally expensive and require specialist expertise. In this study, we demonstrate that graph neural networks trained on coarse-grained MD simulation data can predict phosphoinositide lipid interaction sites on Pleckstrin Homology (PH) domain structures, a large family of membrane binding domains. The predictions are comparable to the results of simulations and require only seconds to compute. Comparison with experimental data shows that the model can predict known phosphoinositide interaction sites and can be used to form hypotheses for PH domains for which there is no experimental data. This model is a next generation tool for predicting protein-lipid interactions of PH domains and offers a basis for further development of models applicable to other membrane protein classes.

## Introduction

The transient association of peripheral membrane proteins (PMPs) with the surface of cell and organelle membranes is a vital physiological process, which plays a critical role in cellular signalling, metabolism, transport, defence and morphology (*1–10*). PMPs are known to bind to membranes through multiple physical mechanisms including non-specific electrostatic interactions, hydrophobic interactions with the membrane core, curvature sensing, cation-π interactions and molecular recognition of particular lipid species from the complex population of lipids present in physiological membranes (*1, 11–18*). Due to their implication in a number of disease processes and their potential for allosteric modulation, drugging the protein-membrane interface of PMPs represents a promising but somewhat under-utilised therapeutic modality (*19–24*).

A major limitation in peripheral membrane protein research lies in the difficulty in experimentally identifying protein-lipid interaction sites and characterizing these in molecular detail (*18, 25–29*). Through structural biology techniques the 3-dimensional atomic structure of PMPs can be determined, but in such structures bound lipids are commonly lost, poorly/partially resolved or do not reveal the lipid identity. Recent advances in cryo-electron microscopy have enabled the resolution of ordered and tightly-bound lipids in some cases, but this is mostly limited to integral rather than peripheral membrane proteins (*26, 30*). A plethora of biophysical techniques allow the measurement of the thermodynamics and kinetics of membrane association and of lipid-binding specificity, but these can be laborious, difficult to perform in a high-throughput fashion and they do not typically reveal all the molecular details of the protein-lipid interactions (*25, 26*). Native mass spectrometry has emerged as a powerful tool for the identification of protein-lipid interaction sites on integral membrane proteins, but to date few studies have applied this technique to peripheral membrane protein-lipid interactions (*31–36*).

Computational methods, particularly molecular dynamics (MD) simulations, can partially mitigate these difficulties by enabling researchers to computationally study membrane protein interactions and dynamics (*11, 37–40*). Notably, MD simulations allow the de novo identification and detailed molecular characterization of protein-lipid interaction sites. Integration of predictions from MD simulations with experimental approaches has had significant impact on our understanding of protein-lipid interactions (*33, 41–49*). The computational cost of MD simulations can be reduced by coarse-grained molecular dynamics (CGMD), which enables the simulation of larger systems for longer timescales although at lower resolution than fully-atomistic systems (*40*). Despite the reduced resolution, CGMD simulations have proven capable of correctly identifying crystallographic lipid binding sites on PMPs and have predicted additional interaction sites which are missing in crystal structures (likely too weak/disordered) but consistent with additional experimental evidence (*50, 51*). The reduced simulation time of CGMD simulations and availability of increasing computational capability has allowed high-throughput simulations of protein-lipid interactions in some cases (*50, 52–55*).

Despite these successes, MD simulations have the limitations of requiring access to high performance computing infrastructure, computational expertise and substantial computation time (even for CGMD simulations), with associated environmental costs. This prevents widespread use of high throughput MD simulations and limits the accessibility of these techniques to a wider community of researchers. There is therefore a need for alternative computational tools which could provide similar insights into membrane biology as MD simulations, but with reduced computational cost and accessibility requirements. Machine learning presents an attractive avenue to build such next generation tools for computational membrane biology, but requires suitable datasets of protein-lipid interactions to train on.

We recently presented a dataset of high-throughput MD simulations of the interactions of 100 pleckstrin homology (PH) domains with a complex plasma membrane model, and demonstrated the capability of the simulations to correctly identify experimentally verified sites of interaction with anionic phosphoinositide lipids, and to provide insights missing from the experimental datasets (*49, 50*). PH domains are the largest family of membrane binding protein domains present in the Pfam database (*56–58*). Comprised, on average, of approximately 140 amino acids, PH domains are defined by their conserved structural motif, consisting of a 7-stranded beta-barrel, capped at one end of the barrel with a C-terminal alpha helix. The other, uncapped end of the barrel typically presents a positively polarised interface for membrane interaction. The variable loop regions between the strands are enriched in basic amino acids which facilitate membrane binding through interaction with anionic lipid headgroups, especially phosphoinositides (PIPs) (*50, 58*). Several PH domains are known for their high affinity binding to phosphoinositide lipids at a so-called canonical site in the open uncapped end of the barrel (*16, 59*). The prevalence of specific phosphoinositide binding properties within the family has been difficult to elucidate, but the most recent investigation concluded that at least half of all human PH domain containing proteins bind specifically to phosphoinositides (*58*). The classical view of PH domain membrane binding, based on the early crystallographic evidence, involves membrane binding driven by high affinity one-to-one binding of a phosphoinositide molecule at the canonical site (*17*). Developments in recent years have unveiled further complexity, including non-canonical phosphoinositide binding sites, the possibility of multiple lipid binding and coincidence sensing, the role of generic interaction with anionic and zwitterionic lipids, and lipid clustering in the recruitment of PH domains to membranes (*47, 50, 51, 60–65*).

In this study we used our previously published MD simulation dataset to train light-weight neural network models utilising a graph attention mechanism to predict the phosphoinositide interaction sites, and to assess their accuracy and speed compared with the results of MD simulations (*50*). Here we show that this new tool allows researchers without access to, or expertise in, MD simulations to rapidly obtain predictions of protein-lipid interactions for their PH domain(s) of interest.

## RESULTS

### Light weight fully-connected graph attention network model to predict phosphoinositide interaction sites on PH domains

Due to the general importance for PH domains of interactions with phosphoinositide (PIP) lipids over all other lipids, we have focussed in this work on predicting PIP interacting sites. The training and validation data consists of CGMD simulations of the membrane binding of 100 PH domains to a complex model of the human plasma membrane inner leaflet. From this dataset we have for every residue of each PH domain annotations of the frequency with which it was in contact with the headgroup of a phosphoinositide (PIP_3_ or PIP_2_) molecule during the final 200 ns of the simulation (see methods). As this contact frequency is convolved with the overall membrane binding affinity of the PH domain, it is convenient for comparison between PH domains to normalize this data by the maximum frequency of contacts exhibited by a single residue in the PH domain, as reported in other simulation studies (*49, 50, 66, 67*). This gives the normalized contact frequency, a number between 0 and 1, representing the relative probability of contact with a phosphoinositide headgroup; the residue with the value of 1 is the residue with the highest probability of interacting with the lipid. The normalized contact frequency observed in the MD simulations provides the reference ground truth data which we aim to train the model to predict. Rather than training the model to predict the normalized contact frequency directly however, we chose to train the model to predict the contact frequency (not normalized), performed normalization outside of the training loop, and assessed the performance between the normalized predicted frequency of contacts and the normalized ground truth data. This decision was based on early empirical evidence that normalization outside of the training loop provided better performance, but without exhaustively testing many possible alternative training settings or network designs.

To predict PH domain/PIP interactions, a graph attention model was developed (Fig. 1). In this model we represent the protein as a fully-connected graph, where each amino acid is a node which is connected to all other aminos acids by an edge for which we include a 4-dimensional feature vector containing the Euclidean distance and the unit vector in the direction α-carbon_A_ −> α-carbon_B_ determined from the protein structure (*68, 69*). At the node level, each amino acid is represented by a 25-dimensional feature vector processed from the sequence and structure. The node feature vectors consists of the one-hot encoding of amino acid identity (a 20-dimensional categorical representation where the position corresponding to the selected amino acid has the value 1, and all other positions have the value 0), the one-hot encoding of DSSP simplified secondary structure class (3 dimensions, from the Dictionary of Secondary Structure of Proteins algorithm), the Shrake-Rupley solvent accessibility (1 dimension) and the total elementary charge within an 8 Å radius of the amino acid α-carbon (1 dimension). Representing the protein as a fully-connected graph of residues, rather than an alternative more sparse connectivity (selected for example by k-nearest neighbours, radial distance or sequence neighbourhood) allows the model to decide for itself which neighbouring or distant amino acids to pay attention to when predicting a residue’s lipid interactions. Although individual lipid interaction sites will be formed by local clusters of amino acids, the effects of more distant amino acids can also be relevant; for example distant lipid interaction sites may either cooperatively enhance or compete with one another, depending on their geometrical relationship with respect to the each other and whether it is possible for both sites to be in contact with the membrane simultaneously. Neighbouring residues can furthermore be highly irrelevant for lipid interaction if they are for example likely to remain buried within the core of the protein and not surface exposed for membrane interaction. An attention mechanism based on GAT-v2 is used to weigh up the relevance (attention score) to each amino acid of the features of all connected aminos acids, taking into considering the amino acid node features and the geometrical information encoded in the edge features (*69*). This enables the model to take into consideration the influence of both local and distant amino acid properties for predicting the lipid interactions of each amino acid. We use two multi-headed graph attention layers, followed by a fully-connected output layer.

**Figure 1.**
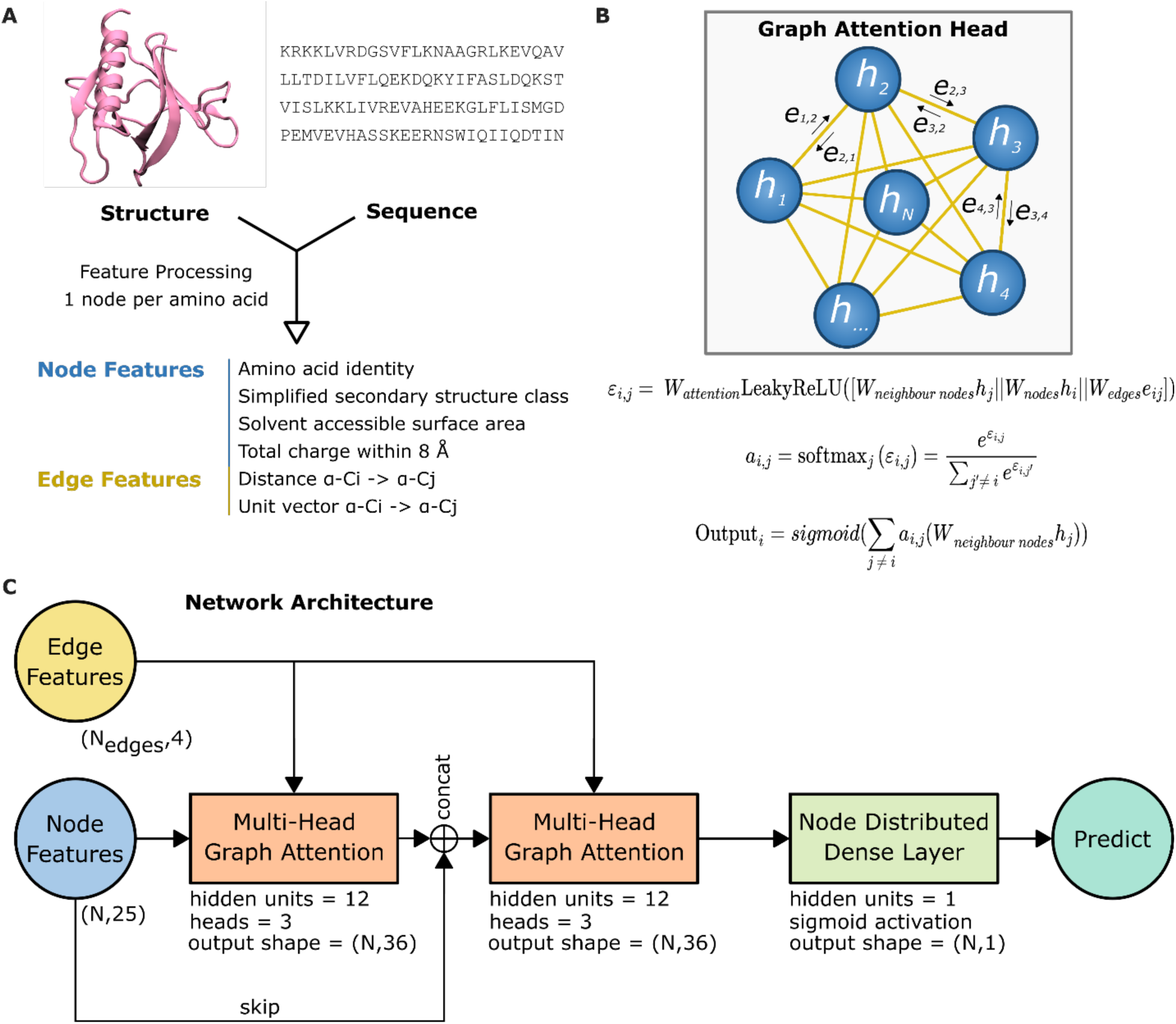
Overview of graph attention network model. **A)** feature processing of the PH domain structure and sequence leads to a graph representation, in which each residue is represented by node features which embed the unique physicochemical properties of that residue. Every residue pair is connected by edges which embed their pairwise geometrical relationship. **B)** Mechanism of a graph attention head based on GAT-v2, which transforms node and edge features into an output. The attention mechanism determines how an amino acid’s lipid interactions are influenced by the properties all other amino acids in the protein. Uppercase Ws represent trainable model parameter matrices. [x||y] represents the operation of concatenating x and y. The attention co-efficient ε*_i,j_* for a target (*i*) and connected (*j*) node is calculated from the node features of *i*, the node features of *j*, and the edge features of the edge *e_i,j_*. The edge attention score for *a_i,j_* is calculated by softmax over all attention coefficients for the target node *i*. The output for the target node *i* is finally calculated as a sigmoid function of the sum of all weighted connected node properties, multiplied by the edge’s attention score. Nodes are not self-connected. **C)** The overall model architecture is comprised of two multi-head graph attention layers (head ouputs are concatenated), with a skip connection, followed by a node distributed dense layer, outputting a per-residue prediction.

The fully-connected graph attention architecture bears resemblance to multi-headed attention transformer models, which have been recently achieving excellent performance in language tasks and other domains. Although transformers and the fully-connected GATv2 model use a different attention mechanism, they are similar in the use of a global attention mechanism over a fully-connected graph, as the sequence inputs of transformers can also be thought of as fully-connected graphs, where the attention mechanism gives weights to the edges. However in contrast to transformers and conventional deep learning architectures we use a much smaller feature and parameter space (akin to classical machine learning methods) to avoid overfitting on the small dataset. By encoding geometrical information, the graph attention mechanism should furthermore allow the model to better account for geometrical relationships between amino acids in a protein compared with a sequence based model (*70–73*). This approach allows us to balance the benefit from the more complex molecular representation available with graph deep neural network architectures with the better performance on small datasets offered by classical machine learning approaches. Further details of model architectures are provided in the Methods and Materials section.

### Model cross-validation

Model performance was assessed using an extensive multiple k-fold cross-validation strategy. In k-fold cross-validation the data is partitioned randomly into k different folds/sets of training and validation data, such that every sample in the dataset is used as a validation sample in only one fold, and as a training sample in all of the (k-1) other folds. A model is trained and evaluated for each training and validation dataset pair (i.e. for each fold). K-fold cross-validation allows better evaluation of model performance than a single train-test fit, but at greater computational cost due to the requirement of training multiple models (*74*). The choice of K is an important hyperparameter, with 10-fold cross-validation being the most commonly used; larger K allows more data for training but means the model is evaluated against a smaller validation set. Furthermore repeating K-fold cross-validation with different randomization seeds results in different combinations of training and testing data in each fold. As our light weight graph attention model is not computationally expensive to train, we have performed multiple k-fold cross validation with k = 5, 10 and 20. For each choice of k we further perform cross-validation using two different random seeds to even more thoroughly evaluate model performance across distinct partitionings of the data. The partitioning of the data for cross-validation is provided in Supplementary Table S1.

Models were assessed on the basis of seven metrics: mean-squared error (MSE), Wasserstein distance (WS distance) and additionally the accuracy, sensitivity, specificity, precision and F1-score (a measure which incorporates both sensitivity and precision) at the task of predicting amino acids with ground truth normalized contacts >= 0.8. MSE calculates the average error per amino acid, and serves also as the training loss function, but does not adequately measure the overall similarity in the shape of the predicted and ground truth distributions, or the capability of the model to identify the top lipid interacting amino acids. The Wasserstein distance provides a measure of overall dissimilarity between the predicted and true distribution (treating these as probability distributions), and can be thought of intuitively as the minimum amount of work (‘earth mover distance’) required to transform the predicted distribution into the ground truth distribution, if the distributions are to be thought of in the Euclidean space as equal mass mounds of dirt. WS distance was selected over other popular distribution metrics such as the Kullback–Leibler (KL) divergence, because unlike KL divergence it is a symmetric metric which is sensitive to locality (i.e. distance in the protein sequence space). Lastly to estimate the capability of the models to detect the most important residues for the protein-lipid interaction, we selected an arbitrary normalized contacts threshold of 0.8 and measured the accuracy, sensitivity, specificity, precision and F1-score in identifying amino acids with normalized contacts >= 0.8, treating this as a binary classification problem. This threshold was selected because we previously found, using the same MD simulation methodology in combination with biophysical experiments to study the peripheral membrane protein PLCγ1, that all amino acids with simulated phosphoinositide normalized contacts >= 0.8 affected the membrane binding affinity of the protein experimentally and that this arbitrary cut-off would also be relevant for PH domains (*49*). These are by no means the only amino acids relevant for membrane binding, but high accuracy and especially sensitivity of predictions at this threshold would make the model a useful tool for identifying phosphoinositide interaction sites. As a baseline, a naive model predicting normalized contacts at random over the whole dataset was empirically found to have MSE = 0.23, mean WS distance = 12, sensitivity = 0.2, precision = 0.06 and F1 score = 0.09.

Summarised cross-validation metrics are provided in Table 1, with more detailed evaluation metrics provided in Supplementary Tables S2–S7. The low variation in all metrics across extensive cross-validation tests indicates that the light weight graph attention architecture is capable of learning robust models which generalize well to unseen samples in the dataset. MSE error ~0.024 indicates strong overall predictive performance. In comparison with the most closely related work to this study, Wang et. al report an overall test set MSE of 0.048 for predicting generic lipid tail contact frequencies of structurally diverse integral membrane proteins (*75*). Predicting phosphoinositide interaction sites on PH domains may be an easier task, due to the strong structural conservation of PH domains and the likely existence of sequence motifs predictive of phosphoinositide interaction (*56*). Sensitivity of ~0.74 at predicting normalized contacts at and above the 0.8 threshold shows good performance in identifying these most likely phosphoinositide interacting amino acids. An F1 score of ~0.69 where the positive class is a minority indicates a good balance between sensitivity and precision. False negatives, which are incorrectly predicted below the 0.8 threshold are still predicted on average at high values below the threshold, on average above the top 90^th^ percentile of predicted values below 0.8.

**Table 1.**
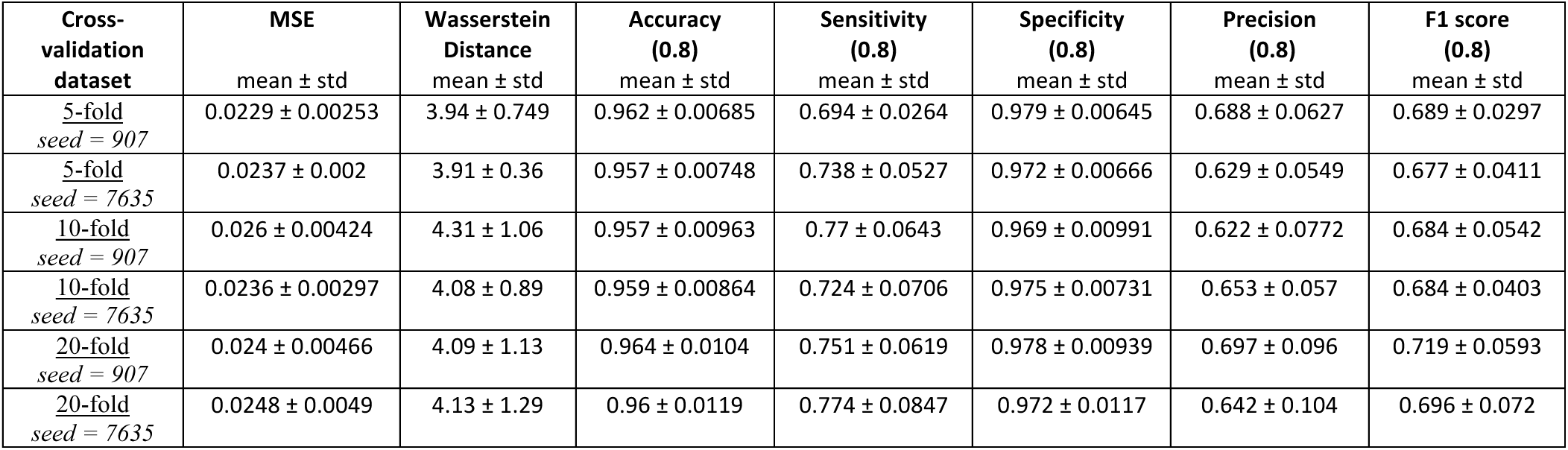
Summary of k-fold cross-validation statistics. 5-fold, 10-fold and 20-fold cross-validation was repeated each with two different randomizations of the PH domain data (random seeds = 907 or 7635). For each cross-validation, metrics were calculated on the validation set in all folds and are reported here as mean ± standard deviation across the folds. Prediction accuracy, sensitivity, specificity and the predicted value of false negatives were calculated by treating the prediction as a binary classification problem, where predicted normalized contacts greater than or equal to a threshold of 0.8 are counted as the positive class; these measures assess the capability of the models at predicting minority group of amino acids with the highest normalized contacts, i.e. those involved in the most probable interaction sites. Detailed statistics are presented in Supplementary Tables S2–S7.

| Cross-validation dataset | MSE<br>mean ± std | Wasserstein Distance<br>mean ± std | Accuracy (0.8)<br>mean ± std | Sensitivity (0.8)<br>mean ± std | Specificity (0.8)<br>mean ± std | Precision (0.8)<br>mean ± std | F1 score (0.8)<br>mean ± std |
| --- | --- | --- | --- | --- | --- | --- | --- |
| <u>5-fold</u><br>seed = 907 | 0.0229 ± 0.00253 | 3.94 ± 0.749 | 0.962 ± 0.00685 | 0.694 ± 0.0264 | 0.979 ± 0.00645 | 0.688 ± 0.0627 | 0.689 ± 0.0297 |
| <u>5-fold</u><br>seed = 7635 | 0.0237 ± 0.002 | 3.91 ± 0.36 | 0.957 ± 0.00748 | 0.738 ± 0.0527 | 0.972 ± 0.00666 | 0.629 ± 0.0549 | 0.677 ± 0.0411 |
| <u>10-fold</u><br>seed = 907 | 0.026 ± 0.00424 | 4.31 ± 1.06 | 0.957 ± 0.00963 | 0.77 ± 0.0643 | 0.969 ± 0.00991 | 0.622 ± 0.0772 | 0.684 ± 0.0542 |
| <u>10-fold</u><br>seed = 7635 | 0.0236 ± 0.00297 | 4.08 ± 0.89 | 0.959 ± 0.00864 | 0.724 ± 0.0706 | 0.975 ± 0.00731 | 0.653 ± 0.057 | 0.684 ± 0.0403 |
| <u>20-fold</u><br>seed = 907 | 0.024 ± 0.00466 | 4.09 ± 1.13 | 0.964 ± 0.0104 | 0.751 ± 0.0619 | 0.978 ± 0.00939 | 0.697 ± 0.096 | 0.719 ± 0.0593 |
| <u>20-fold</u><br>seed = 7635 | 0.0248 ± 0.0049 | 4.13 ± 1.29 | 0.96 ± 0.0119 | 0.774 ± 0.0847 | 0.972 ± 0.0117 | 0.642 ± 0.104 | 0.696 ± 0.072 |

### Further examination of model performance

Our model provides excellent predictions for the vast majority of the PH domains that we have examined, but there are some cases in which there are greater limitations to its accuracy. The best performing results from the plcd1, IQSEC1, TBC1D2 and ARAP2 PH domains (Fig. 2A-D) exhibit remarkable performance, providing highly comparable predictions as the MD simulation with only small differences discernible by inspection. This can be seen in detail for the plcd1 PH domain visualized on the crystal structure in Figure 3A. The model excellently predicts the key phosphoinositide interacting residues around the β1 and β2 strands, the β1-β2 loop and the β3-β4 loop, directly comparable to the MD simulation. The best and worst (on the basis of MSE) validation set predictions for all PH domains are provided in supplementary figures S2–S8.

**Figure 2.**
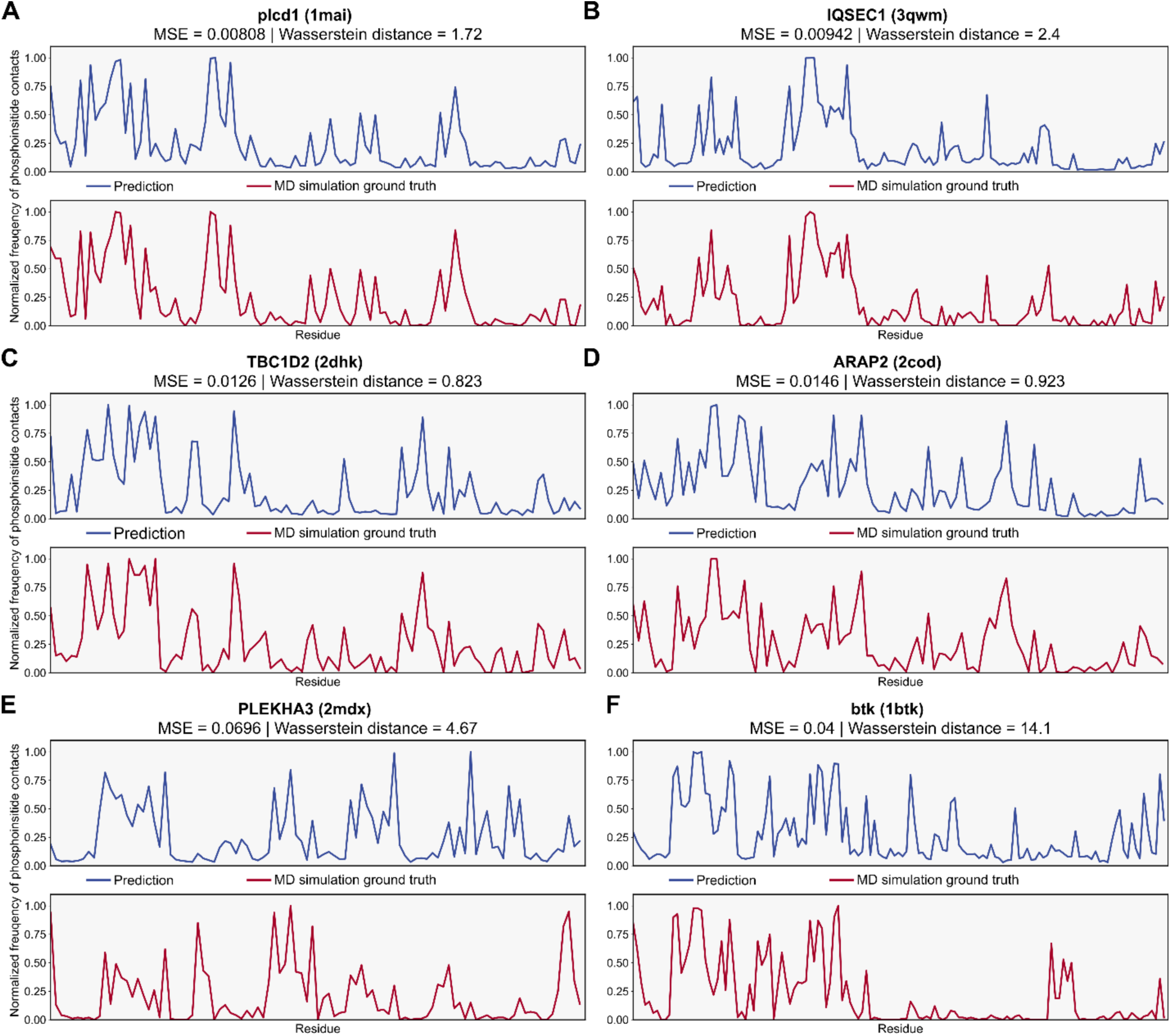
Best (A-D) and worst (E-F) predictions across all validation samples. Normalized frequency of contacts with phosphoinositide headgroups predicted by the graph attention model (blue), versus the ground truth MD simulation data (red) shown for a selection of PH domains with the top two best and the worst predictions, quantified by mean-squared error **(A, B, E)** or Wasserstein distance **(C, D, F)**. The PH domain gene name is displayed above each plot, with the PDB ID code in brackets.

**Figure 3.**
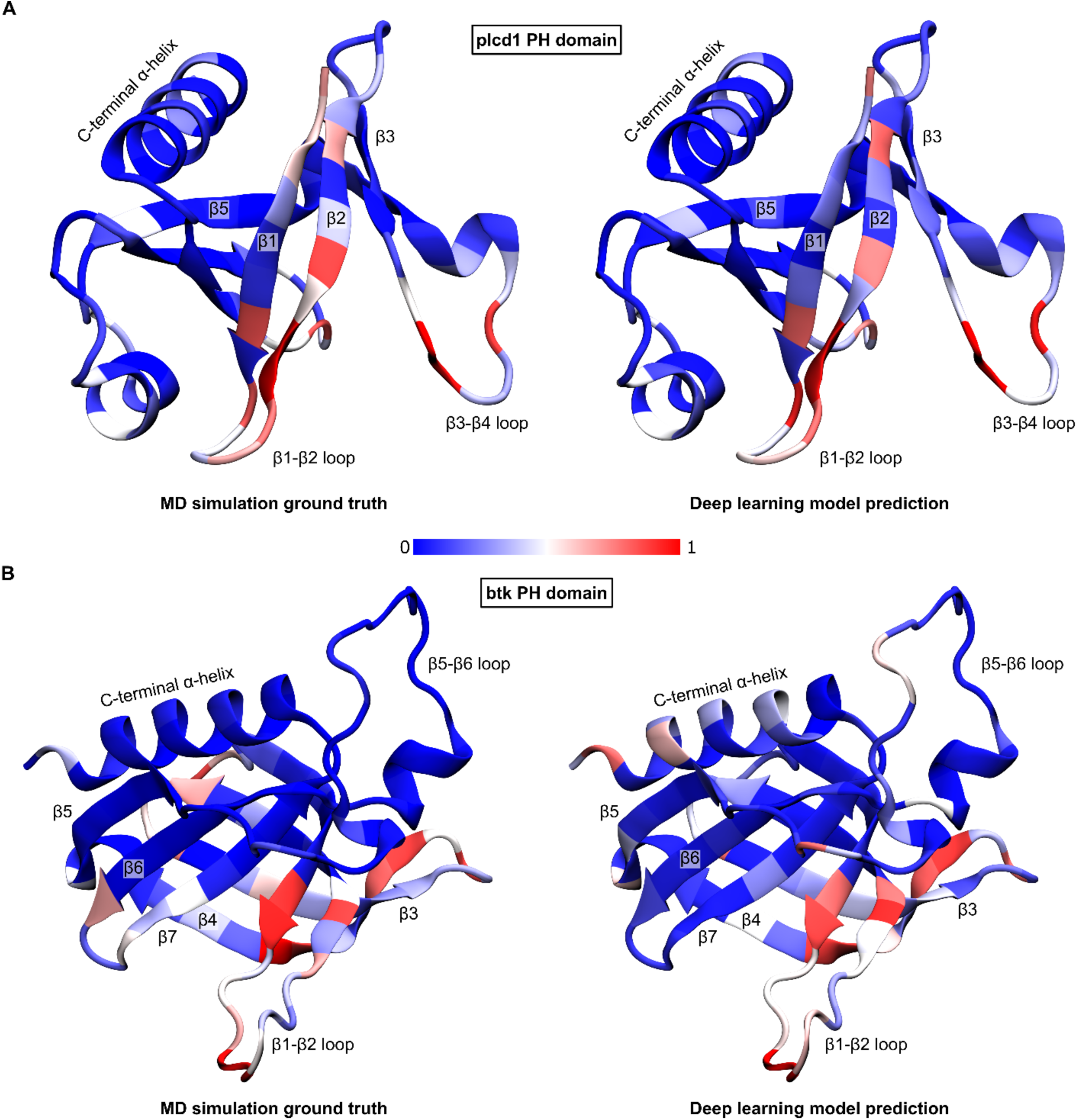
Structural visualization of normalized phosphoinositide headgroup contact frequencies by MD simulation and deep learning model prediction. Normalized frequency of contacts with phosphoinositide headgroups for **A)** plcd1, the PH domain with the lowest MSE observed during cross validation (MSE = 0.00808, WS distance = 1.72) and **B)** btk, the PH domain with the highest WS distance observed during cross validation (MSE = 0.04, WS distance= 14.1). Reference MD simulations (left) are compared with the deep learning prediction (right). Normalized frequency of contacts is coloured per residue on a scale from 0 to 1. Relevant secondary structural elements are labelled.

Even though the results from the PLEKHA3 and btk PH domains are worst performing (Fig. 2E-F) both show generally good prediction of the lipid interaction sites but with some inaccuracies, where the model has overestimated the importance of some binding sites while underestimating others. The prediction for the btk PH domain (Fig. 2F) performs well for the first half of the sequence containing the main phosphoinositide interacting residues, but the model has inaccuracies in the second half of the sequence – it suggests false interactions in the β5-β6 loop and C-terminal α-helix, and fails to identify moderate interactions contributing to the canonical site by tyrosine, glycine and leucine residues on the β6 and β7 loops. A more detailed comparison of the ML prediction and the reference MD simulation is visualised on the protein structure in Figure 3B.

The second half of the β5-β6 loop contains four lysine and arginine residues, which normally when present in this loop region form a non-canonical binding site as in the Arhgap9 PH domain (*50*). There is additionally a positively charged ridge along the C-terminal α-helix aligned along the same side of the β-barrel as these lysines and arginines in the β5-β6 loop; these are likely the cause for the model’s prediction of moderate interactions at these locations. However, the unusually extended β5-β6 loop of the btk PH domain contains 7 negatively charged glutamic acid and aspartic acid amino acids, which in the simulation neutralises the effect of the basic residues and repels this region away from the membrane, causing this PH domain to adopt a more upright orientation of the β-barrel on membrane rather than the side-on orientation adopted by PH domains such as Arhgap9 and as predicted by the ML model (*50*). The model has in this case slightly erred in understanding how the global electrostatic properties will affect the overall orientation of the PH domain on the membrane. However, when examining the residues with the highest interaction, around the β1, β2, β3 and β4 loops, the model makes good predictions in reference to the MD simulation. Therefore despite being the worst from all the cross-validation datasets in terms of Wasserstein distance, the prediction is still quite meritorious in this regard, and would be useful (despite its deficiencies) to membrane biologists looking to gain computational insights into phosphoinositide interacting residues of the btk PH domain without running MD simulations.

### Evaluating predictions on out-of-dataset PH domain structures and comparison with experimental observables

To further assess the generalisability and practical applicability of the learned graph attention models it is fruitful to make predictions for PH domain structures not contained in the MD simulation dataset and for which structural data is available for the phosphoinositide binding sites. As the MD simulation dataset comprises almost all non-redundant human, rat and mouse PH domain structures published up to 2022, there are few PH domains suitable for this further analysis. Fortunately there is the human Arap3 PH1 domain structure solved in complex with the short-tailed phosphoinositide analogue diC4-PI(3,4,5)P_3_, published in 2023 by Zhang et al., with additional NMR chemical shift perturbation (CSP) experiments (*76*). Their data demonstrate specific recognition of diC4-PI(3,4,5)P_3_ in the canonical site, with key interactions with the headgroup stabilised by K296, R308, K329 and R355 and supplemented by S298, Q306 and Y319. Fig. 4A shows the predictions of a trained graph attention model for this structure (PDB ID: 7YIS), along with the position of the complexed diC4-PI(3,4,5)P_3_ observed in the structure. Residues predicted by the model to have normalized frequency of contacts >= 0.8 are displayed as blue sticks and labelled. This analysis excellently predicts all basic residues which interact with the crystallised diC4-PI(3,4,5)P_3_ in the canonical pocket. Outside the canonical site the model predicts six additional residues (R307, R341, K344, K347, R360 and R382) as additional residues with high probability of phosphoinositide headgroup interaction. The geometrical arrangement of these residues is striking, as they form an electropositive ridge along the top of the putative membrane-binding interface. It is highly plausible that these residues could enhance the overall membrane-binding affinity through additional interactions with phosphoinositides and other anionic lipids at the membrane interface, supplementing primary binding at the canonical site (*50, 51, 65*). ^1^H-^15^N HSQC NMR experiments indeed show chemical shift perturbation of R307, K347 and R360 above the mean upon addition of soluble diC4-PI(3,4,5)P_3_, with R341 perturbation just below the mean and the residue K344 not resolved (*76*). These perturbations were attributed to conformational change upon ligand binding to the canonical site, however interactions with diC4-PI(3,4,5)P_3_ could also have this effect. Note that these experiments were performed with soluble phosphoinositide analogues and not with binding to a membrane model; it may be the case that the additional interactions outside of the canonical site could be dependent on the presence of the membrane interface. There was no perturbation at R382. The additional interactions, although plausible, may be false positives; nevertheless the model clearly demonstrates generalisability and practical usefulness by identifying the known locations of phosphoinositide interactions site and suggesting additional plausible sites which warrant further experimental investigations.

**Figure 4.**
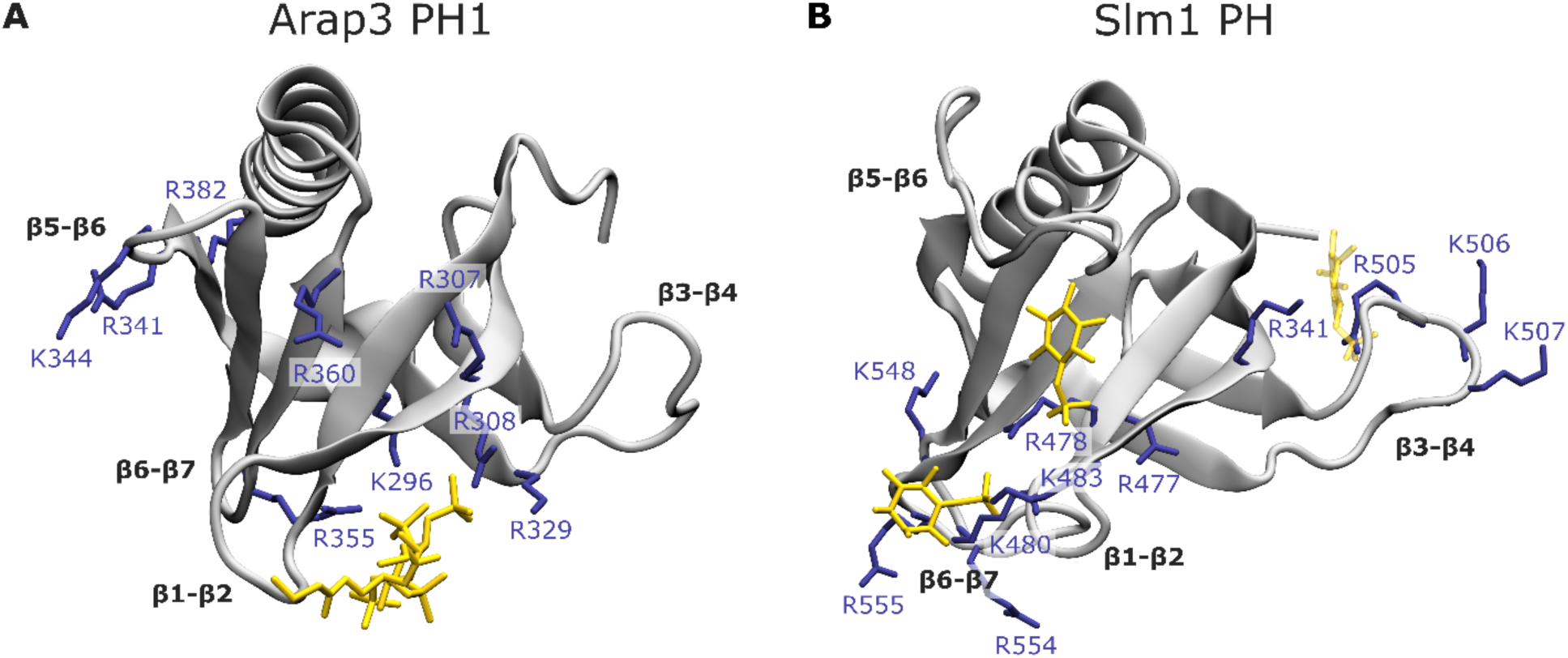
Comparison of predictions with structural data for the Arap3 and Slm1 PH domains. Structures of **A)** human Arap3 PH1 domain (PDB ID: 7YIS) in complex with diC4-PI(3,4,5)P_3_ and **B)** yeast Slm1 PH domain in complex with inositol 4-phosphate. Amino acids predicted by the deep learning model to have normalized frequency of contacts >= 0.8 are displayed as blue sticks and labelled. Co-crystallised phosphoinositide analogues and partially resolved phosphate are displayed as yellow sticks. Relevant unstructured loop regions between beta-sheets are labelled. In panel B the unliganded 4A5K protein structure is displayed and used for the prediction, as other structures were missing residues in the β5-β6 loop. STAMP structural alignment was performed between the 4A5K structure and the ligand containing 4A6K (RMSD = 0.805 Å) and 4A6H (RMSD = 0.530 Å) structures, and the inositol 4-phosphate shown are from the these aligned structures. The structural alignments are presented in Supplementary Figure S9 and the predictions for all amino acids (not just those >= 0.8) are visualised in Supplementary Figure S10.

Similar realistic predictions are obtained for the yeast Slm1 PH domain (Fig. 4B), which is not in the training dataset as this is comprised only of mammalian PH domains. Crystal structures of the Slm1 PH domain in complex with inositol-4-phosphate resolve multiple non-canonical sites of phosphoinositol interaction, particularly at a pocket formed between R478 and Y485 as well at sites around the β6-β7 and β3-β4 loops (*60*). Impressively the model predicts a high frequency of phosphoinositide interactions around all of the crystallographically observed sites. The model additionally proposes the existence of interactions at a more canonical-like site formed at R477. Binding is not observed here in the crystal structure, but mutagenesis studies probing putative lipid binding pockets of Slm1 PH found that the mutation R477A destabilized the membrane association of Slm1 PH at the eisosome (*77*). The graph attention model therefore shares the power of MD simulations to predict important phosphoinositide interaction sites which are absent in crystal structures.

### Application to PH domains of unknown structure

Considering the highly conserved secondary structure pattern of PH domains, with only short disordered regions, AlphaFold predictions of the structures of PH domains may be expected to be quite reliable (*78*). The combination of AlphaFold with our graph attention model could extend the predictions of phosphoinositide interaction sites to PH domains of unknown structure. We demonstrate this capability for three example PH domains – the DGKH PH domain, the RASA3 PH domain and the PH domain of the VAN3-binding protein from Mexican cotton (Gossypium hirsutum).

Diacylglycerol kinase eta (DGKH) is a member of a family of enzymes which phosphorylate the lipid diacyl glycerol (DAG) to phosphatidic acid. The structure of DGKH has not been solved, but it has been found to contain a PH domain with similar affinity and selectivity for PI(4,5)P_2_ as the plcd1 PH domain (a common high affinity PI(4,5)P_2_ probe) (*79*). It has greater affinity for PI(4,5)P_2_ than other diacylglycerol kinase isozymes. K74A and R85A mutants were found to be unable to bind to the PI(4,5)P_2_. Prediction of the graph attention model using the AlphaFold DGKH PH structure (Fig. 5A) indeed proposes K74 and R85 as highly likely sites of phosphoinositide interaction, in addition to several other basic residues lining the putative membrane interface.

**Figure 5.**
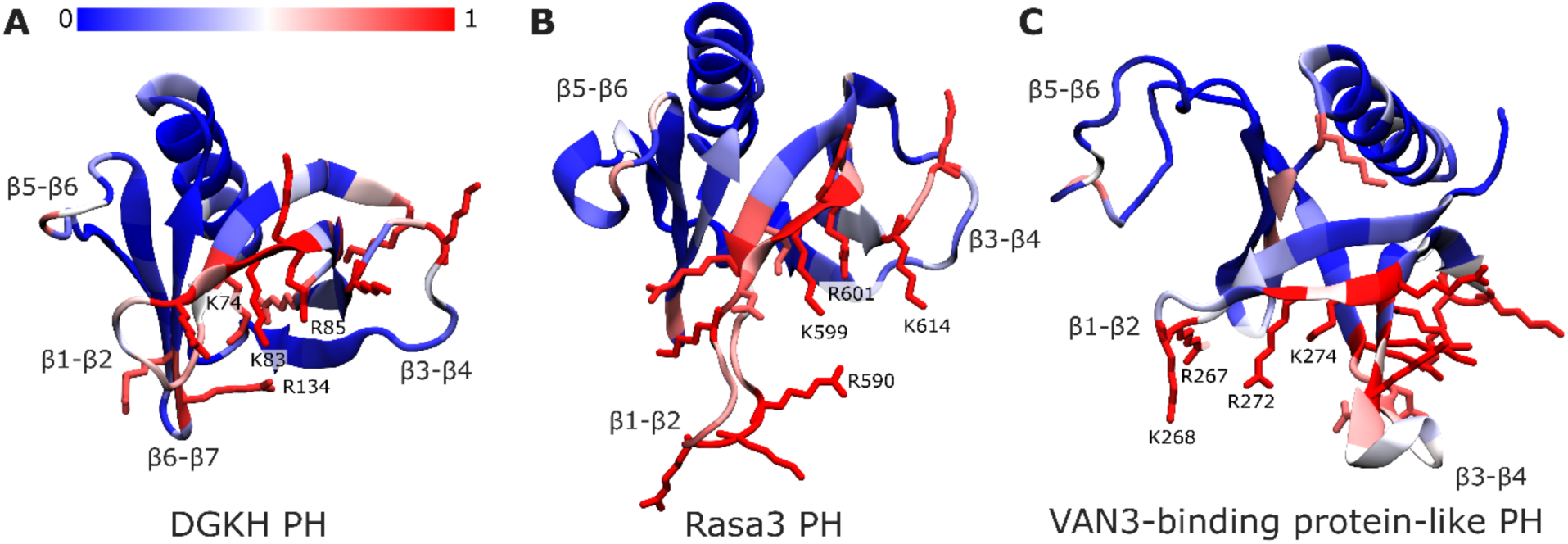
Predicted normalized contact frequency with phosphoinositide headgroups for AlphaFold modelled structures. Structures of **A)** human DGKH PH domain (AF: Q86XP1), **B)** rat Rasa3 PH (AF: Q9QYJ2) and **C)** mexican cotton VAN3-binding protein-like PH (AF: A0A1U8N485). The predicted normalized contact frequency is coloured by residue on the structure as indicated by the colour bar. Residues with predicted normalized contact frequency >= 0.8 are shown as sticks.

The Ras GTPase activating protein 3 (RASA3) is a PH domain containing protein of unknown structure, targeting PI(4,5)P_2_ at the plasma membrane, and has been identified as potential tumour suppressor and possible drug target in breast cancer chemotherapy (*80–83*). Site-directed-mutagenesis found that K585, A587, N597, F598 and R601 are residues important for membrane association (*82*). The graph attention model (Fig. 5B) correctly identifies the basic residues K585 and R601 located near the canonical pocket and highlights additional lysine and arginine residues adjacent to this pocket and on the β1-β2 and β3-β4 loops.

Exploring PH domains in other kingdoms of life, the recently discovered VAN3-binding protein of Gossypium hirsutum contains a PH domain (*84*). Smith et al. have explored the potential phosphoinositide binding of this PH domain by structural alignment and comparison of the AlphaFold model with known PH domains such as btk (*84*). From their alignment they proposed Q264, R272, K274, N303, and L342 as putative phosphoinositide interacting residues. Our model concurs with their analysis, predicting K274 and R272 as strongly interacting residues in the canonical site, alongside H294 and R267. Interestingly the model also highlights seven lysines and arginines oriented along the outward face of the β3 and β4 strands and their connecting loop. This is an usual feature among PH domains and in the absence of competing interactions at this interface (such as protein-protein interactions at the membrane) this highly positively charged face of the PH domain could direct its orientation on the membrane, causing it to adopt a side-on orientation similar to that observed in simulations of the Exoc8 PH domain (*50*).

These examples demonstrate the practical application of our predictive model to PH domains of unknown structure and newly discovered PH domains. This can enable researchers to quickly get useful predictions of phosphoinositide interacting sites which can inform models of membrane interaction, guide experimental design and provide testable hypothesis.

### Prediction beyond PH domains

Despite the fact that the model has been trained using only data for PH domains, we also investigated its generalizability to PMPs which do not belong to the PH domain family. Phox Homology (PX) domains are another structurally conserved family of peripheral membrane protein domains of similar size to PH domains, and which bind to phosphoinositides (*85*).

To examine the generalizability of the model on a PX domain, we selected the p40 PX domain as it was the first identified PX domain, a crystal structure (PDB ID: 1h6h) in complex with phosphatidylinositol 3-phosphate is available, and there is further additional MD simulation data for the membrane association of p40 PX reported in the literature (*86, 87*). The crystal structure has a conserved phosphoinositide binding site formed by R58, R60, K92 and R105 (Fig. 6A) (*86*). MD simulations have furthermore identified a secondary binding site for phosphoinositide interaction facilitated by K32 and R33 in the β1-β2 loop (*87*). All of these residues are presented in red in Figure. 6A. The prediction of the graph attention model is presented for comparison in Figure 6B, where residues predictive to have normalized frequencies of phosphoinositide contacts >= 0.8 are also shown as red sticks. To our immense satisfaction, the model correctly identifies the key residues R58, R60, K92 (but not R105) of the crystallographic binding site, as well as the additional residues K32 and R33 in the β1-β2 region. The model predicts additional residues K46 and K52 as having high frequencies of phosphoinositide interactions but this is not supported by the MD simulations (*87*). The model is therefore making some surprisingly realistic inferences for this PX domain despite being trained only on PH domain data. We do not recommend using the model to make inferences for non-PH domains, as the generalization capability for such proteins has not been quantitatively and systematically evaluated and has been investigated here only for one protein. The results for p40 phox are however a promising indication that further generalization of the model could be achievable, perhaps by further training with an expanded dataset including other types of peripheral membrane proteins.

**Figure 6.**
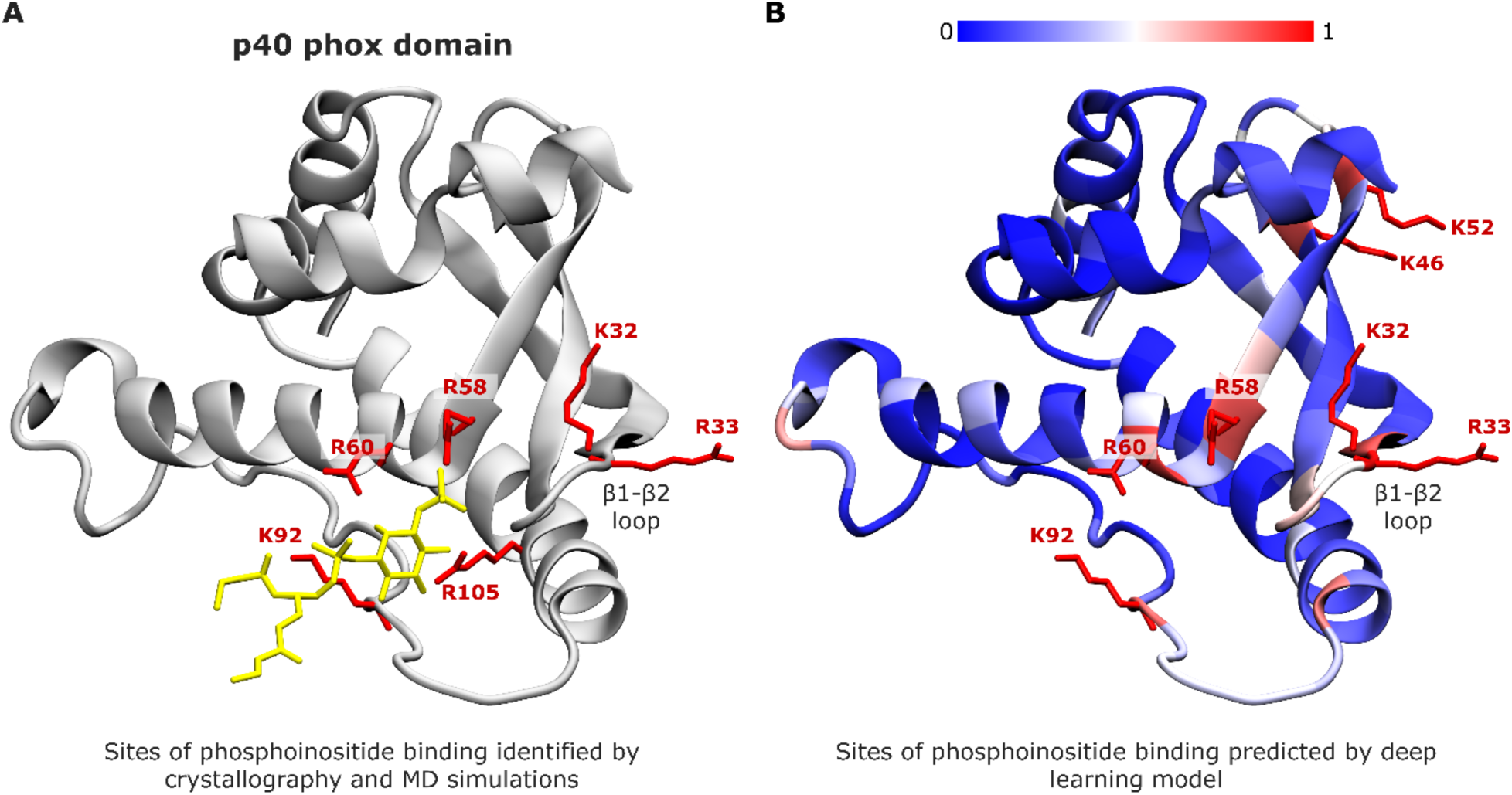
Prediction beyond PH domains: p40 phox. Crystal structure (PDB ID: 1h6h) of the p40 phox domain. **A)** The structure is shown (silver) in complex with phosphatidylinositol 3-phosphate (yellow sticks). Residues which interact with phosphoinositides are shown as red sticks. These residues were selected because they either engage with the bound lipid in the crystal structure (R58, R60, K92, R105) or form a secondary interaction site identified by MD simulations (K32 and R33) (*86, 87*). **B)** The protein structure as in **A**, but coloured according to normalized frequency of contacts with phosphoinositide headgroups predicted by the deep learning model (coloured according to the colour scale). Residues with predicted contacts >= 0.8 are shown as red sticks.

### The light weight graph attention model allows rapid predictions

To estimate the time required for predictions we benchmarked the time for feature processing and inference without GPU acceleration, with a batch size of 1, using a consumer laptop with an 8 core AMD Ryzen 7 6800HS CPU, 16 GB of RAM and running Windows 11. Feature processing and inference were conducted for 100 PH domains. Import of python modules and the saved tensorflow model required ~16 seconds. Feature extraction and inference for the first two PH domains in the dataset required ~10 seconds and ~3 seconds respectively. All subsequent inferences required between 0.3 and 1 second for feature processing and inference. This data is presented in Figure 7. The slower time for the first two inferences can be attributed to a ‘warming up’ period of Tensorflow, arising from processes such as memory allocation and computation caching. The total time required to obtain inferences from 100 PH domain structures was ~80 seconds. This is incredibly fast performance on consumer grade hardware. For comparison, the underlying CGMD simulation data for each PH domain required 20 x ~23 hour simulations using 24 cores on high performance computing infrastructure to obtain convergence (approximately 46,000 hours of supercomputing time for the full dataset). The graph attention model therefore allows prediction of the normalized frequency of contacts with phosphoinositide headgroups for this dataset more than 6 orders of magnitude faster than the MD simulations and with an overall MSE of approximately ~0.024 relative to the simulations.

**Figure 7.**
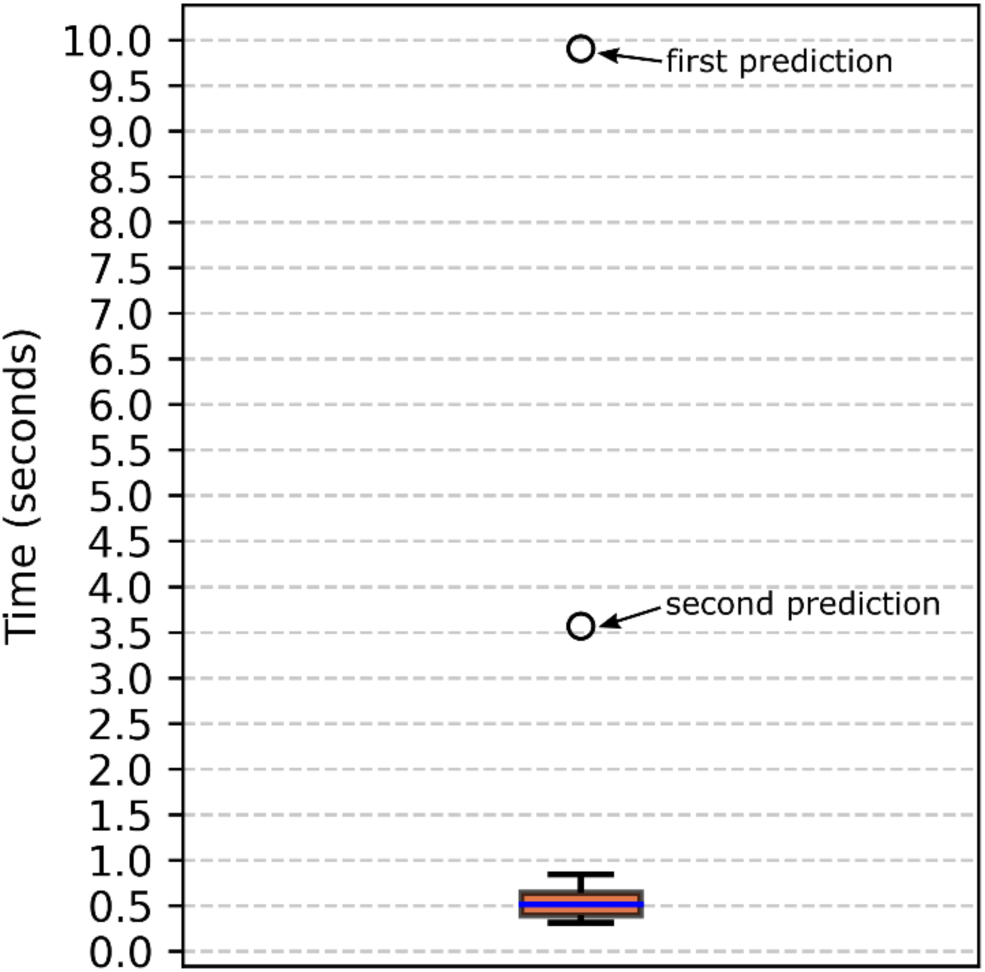
Time required for feature processing and inference for each PH domain. The first two predictions require several seconds due to a ‘warming up’ period of TensorFlow. All subsequent predictions require less than one second to compute. Data does not include the time required for initial loading of python modules and the tensorflow model.

## DISCUSSION

We have presented a graph neural network model for fast prediction of phosphoinositide lipid interaction sites on PH domains, which was trained on high throughput CGMD simulation data. The model exhibits excellent and robust performance across an extensive cross-validation assessment, with MSE of ~0.024 relative to the ground truth MD simulations. The model provides comparable predictions of phosphoinositide interacting residues as CGMD simulations but at many orders of magnitude less computational cost. Further assessment using out of dataset PH domain structures and experimental data indicates good generalization to other PH domains, and this can be extended through the use of AlphaFold to PH domains of unknown structure.

The capability to rapidly predict sites of interactions of specific lipid types with peripheral membrane proteins is highly exciting on two fronts. Firstly, the high compute demand, computational expertise and hardware requirements limit the accessibility of molecular dynamics predictions to the more general community of membrane biology researchers. Our model for PH domains can be run in seconds on standard laptops and PCs, with minimal programming knowledge and software requirements. This makes the useful insights that can be attained by MD simulations easier to achieve and accessible to a wider community. Secondly, the rapid inference speed of our model opens up the possibility for prediction throughput unimaginable using MD simulations (but at reduced accuracy). Considering the importance of peripheral membrane proteins in cellular signalling and disease-linked processes, and the potential for drugging the protein-membrane interface, such a tool could be highly impactful for membrane biology research (*19, 20, 24*). We note that the goal of this research is not to replace MD simulations (which we consider to be a method of superior accuracy and rigour), but to provide a substantially faster and more accessible tool that can provide similar predictions.

Recently, a few pioneering studies have explored the use of machine Learning (ML) for modelling membrane biology. Wang et al. trained a deep convolutional recurrent neural network (DCRNN), a model previously shown to perform well for secondary structure prediction from sequence, on the MemProtMD database of integral membranes protein simulations with a generic DPPC model bilayer (*54, 75*). It was demonstrated that the DCRNN model could predict the contact probability (probability of coarse-grained particles < 6 Å during the final 800 ns of simulation) between lipid acyl tails and each residue of the protein with a good accuracy (MSE = 0.048 on test set) when trained on 5000 membrane protein sequences from MemProtMD. Furthermore, incorporation of the membrane contact probability prediction was shown to improve integral membrane protein structural modelling (*75*). In the case of peripheral membrane proteins, Chatzigoulas and Cournia recently assembled from the available experimental literature a dataset of 54 PMPs with known membrane-penetrating amino acids, and trained ensemble based classical machine learning classifiers to predict the membrane-penetrating residues (*88*). Van Hilten et al. using a convolutional neural network and a dataset of 50000 CGMD simulations, were able to predict the membrane curvature sensing ability of short (7-24 amino acids) peptides with comparable accuracy to the MD simulations (*89*).

Nevertheless, the prediction of interaction sites for specific types of lipids found in complex physiological bilayers using machine learning has not yet been explored, likely due to the lack of datasets of such interactions. Despite being larger than other datasets used for modelling peripheral membrane proteins with machine learning, this 100 PH domain dataset is small from the point of view of machine learning, and 50-fold smaller than the dataset of integral membrane protein simulations used in the work of Wang et al.; avoiding overfitting to the small dataset was therefore a major consideration in our model design (*75, 88*). Developing deep learning architectures which are effective for small datasets is an on-going problem in the field of deep learning research. We focussed especially on developing a light weight model architecture which can simultaneously harness the power of deep learning for learning complex relationships between amino acids while maintaining a small parameter size to avoid overfitting. We employed graph neural networks as these are well suited for representation of molecular structure, and selected a small feature space representing a small number of properties that we thought would be most relevant for predicting lipid interaction properties. We furthermore kept a small parameter size by using fewer hidden units per layer than normally employed in deep learning. These neural network design choices allowed us to benefit from the power of deep learning without overfitting the small dataset and also result in the astonishing prediction speed.

During this work PH domains have been modelled as individual and isolated protein structures, but physiologically they are very often in fact a small domain within the context of a much larger protein sequence, which may contain additional membrane binding domains. The model performance has not been evaluated for predictions on structures in which the PH domain is embedded in a larger protein or complex, and thus is recommended only for use with isolated PH domain structures. This is not a substantial deficiency, as the experimental and computational study of PH domains as isolated structures is a common practice in this research field and has provided great insights into the biochemistry of PH domains and the proteins in which they are embedded.

Model performance has been critically evaluated. Although the overall performance is good, in some cases, such as for the btk PH domain, it makes some mistakes with respect to how the overall orientation of the PH domain on the membrane is influenced by global electrostatics. We therefore recommend that the overall PH domain electrostatics be taken into consideration when interpreting model predictions. In some cases the model under- or over-estimates key interaction sites compared with the MD simulation, as can be seen in Figures 2–3 and Supplementary Figures S1-S8. Predictions are furthermore very likely to be limited by the same fundamental limitations of the underlying CGMD simulation data. This includes inadequate modelling of specificity between different phosphoinositide species/stereochemistries, limited modelling of protein conformational changes, no insight into the overall membrane binding affinity of the PH domain and the absence of consideration of intermolecular protein-protein interactions or the greater protein context of the PH domains. For further details of the merits and limitations of the underlying training data please refer to the publication of this simulation dataset (*50*).

Having considered these limitations, our model provides a practical and accessible tool for obtaining fast insights into putative phosphoinositide interaction sites of pleckstrin homology domains, similar to MD simulations, at substantially reduced computational cost. Considering the difficulties and challenges in experimentally studying membrane and lipid association, our approach presents a method which can help to identify sites of lipid interaction that have been overlooked in the available data and provide hypotheses which can be experimentally tested. Predictions of the model during this study already raise interesting questions, for example about the proposed membrane binding role of the predicted electropositive ridge in the Arap3 PH1 (Fig. 4A). Assessment of the generalization capability of the model beyond PH domains showed promising results in the case of the p40 Phox domain. Expansion of the training MD simulation dataset to include other peripheral membrane protein families and peripheral membrane proteins which do not belong to a domain family, as well as the inclusion of other lipid types are promising directions for future development.

## MATERIALS AND METHODS

All model training and testing was performed with Tensorflow Keras 2.13.0 and python 3.9.17 (*90*).

### Dataset selection and feature processing

The dataset consists of 100 PH domain PDB structures and the results of coarse-grained molecular dynamics simulations as previously published (*50*). In these MD simulations the coordinates of all particles were saved every 0.4 ns. For every amino acid *i* in the protein and for every frame f during the final 200 ns of the MD simulation, 1 contact was counted when any particle of the amino acid was within 5.5 Å of any headgroup particle of a PIP_2_ or PIP_3_ lipid. Contacts were totalled over all 20 simulation replicates for each protein and over all frames for the final 200 ns of the simulation. Let c*_i_* be the total contacts counted for an amino acid *i* in a protein of sequence length |*N|*. Then the contact frequency is equal to *c_i_* divided by the total number of frames during the 200 ns window.

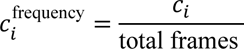

When comparing amino acid contact frequencies between different PH domains, the contact frequencies will be affected also by the differences in overall membrane binding affinity of the PH domain. The simulation methodology however, designed for identifying key lipid interaction sites, does not provide a good quantitative estimate of the overall PH domain-membrane binding free energy. It is therefore common practice to instead examine the normalized contact frequency, where the contacts are normalized relative to the amino acid with the highest number of contacts for the protein (*49, 50, 66, 67*):

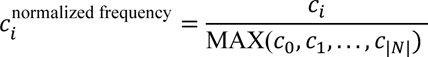

We wish to predict the normalized contact frequency for all amino acids in the protein, however in practice training to predict contact frequency and then normalizing after training was found to be more effective. Models were therefore trained to predict 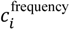 and predictions evaluated on 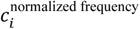, with normalization conducted after training.

### Graph Attention Network Architecture

The model architecture is based on a modification of the GATv2 attention mechanism to include edge features (as shown in Figure 1), and operates on the protein as a fully connected graph (*69*).

### Cross-validation and training

Shuffling of data for 5-fold, 10-fold and 20-fold validation was performed using the model_selection.KFold() class of scikitlearn. For every fold cross-validation was repeated with two different random seeds (seeds 907 or 7635). Models were trained for a maximum of 40 epochs with early stopping, using the Adam optimizer and with mean squared error as the loss function (*94*). The learning rate was 0.0071. For validation metrics all prediction and ground truth contacts frequencies were first normalized to normalized contacts frequencies by all dividing all elements in the tensor by the maximum value of any element in the tensor. Mean squared error (MSE) was calculated using Keras (*90*). Wasserstein distance (WS) was calculated using SciPy, treating the normalized contacts as a probability distribution (*95*). Accuracy, sensitivity and specificity, precision and F1 score at the 0.8 normalized contact frequency threshold were calculated as a binary classification problem, with predictions >= 0.8 counted as true or otherwise false.

### Application of trained model for prediction on out-of-dataset PH domains and other proteins

To make further predictions we use the model trained by fold 7 of the 10-fold cross-validation with seed 907. This was selected arbitrarily, with the reasoning that the 10-fold cross-validation datasets provided the best balance between the number of samples in the training dataset (better generalisability) and the number of samples in the validation dataset (more robust performance assessment); the fold 7 seed 907 model then had the highest F1 score (0.755) of all models assessed during 10-fold cross-validation.

Inferences were made for the Arap3 PH domain (PDB ID: 7YIS), Slm1 PH domain (PDB IDs: 4A5K, 4A6K, 4A6H) and p40 phox (PDB ID: 1h6h) downloaded from the protein databank. Inferences were made for AlphaFold models of the human DGKH PH domain (AF: Q86XP1), rat Rasa3 PH domain (AF: Q9QYJ2) and mexican cotton VAN3-binding protein-like PH domain (AF: A0A1U8N485) after downloading the structure from the AlphaFold Protein Structure Database and extracting the PH domain residues from the larger protein structure as previously described (*50, 78*).

## Funding

This work was funded by the Biotechnology and Biological Sciences Research Council grant BB/M011151/1. This work was undertaken on ARC3 and ARC4, part of the High Performance Computing facilities at the University of Leeds, United Kingdom.

## Author contributions

Conceptualization: A.C.K., F.S., H.W., and K.I.P.L.H.

Methodology: A.C.K., H.W., and K.I.P.L.H.

Acquisition and analysis of data: K.I.P.L.H.

Visualization: K.I.P.L.H.

Funding Acquisition: A.C.K., F.S., and H.W.

Supervision: A.C.K., F.S., and H.W.

Writing—original draft: KI.P.L.H.

Writing—review and editing: A.C.K., F.S., H.W., and K.I.P.L.H.

## Competing interests

The authors declare that they have no competing interests.

## Data and materials availability

All data needed to evaluate the conclusions in the paper are present in the paper and/or the Supplementary Materials. The codebase has been submitted for peer review and will be freely accessible upon publication.

## SUPPLEMENTARY MATERIAL

**Supplementary Table 1.**
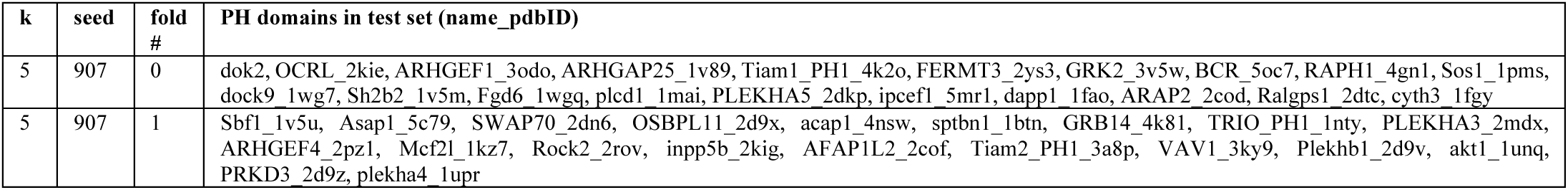
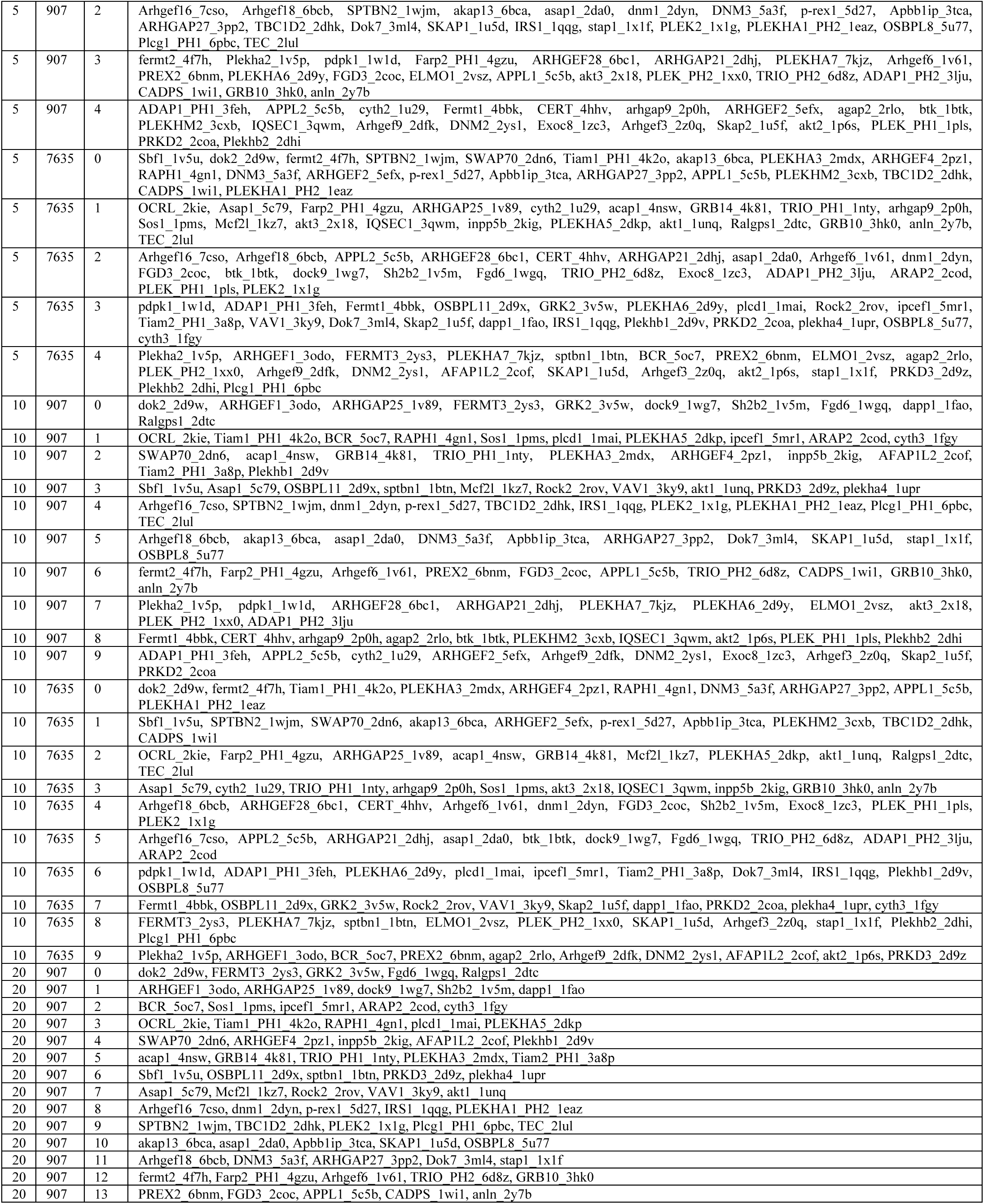
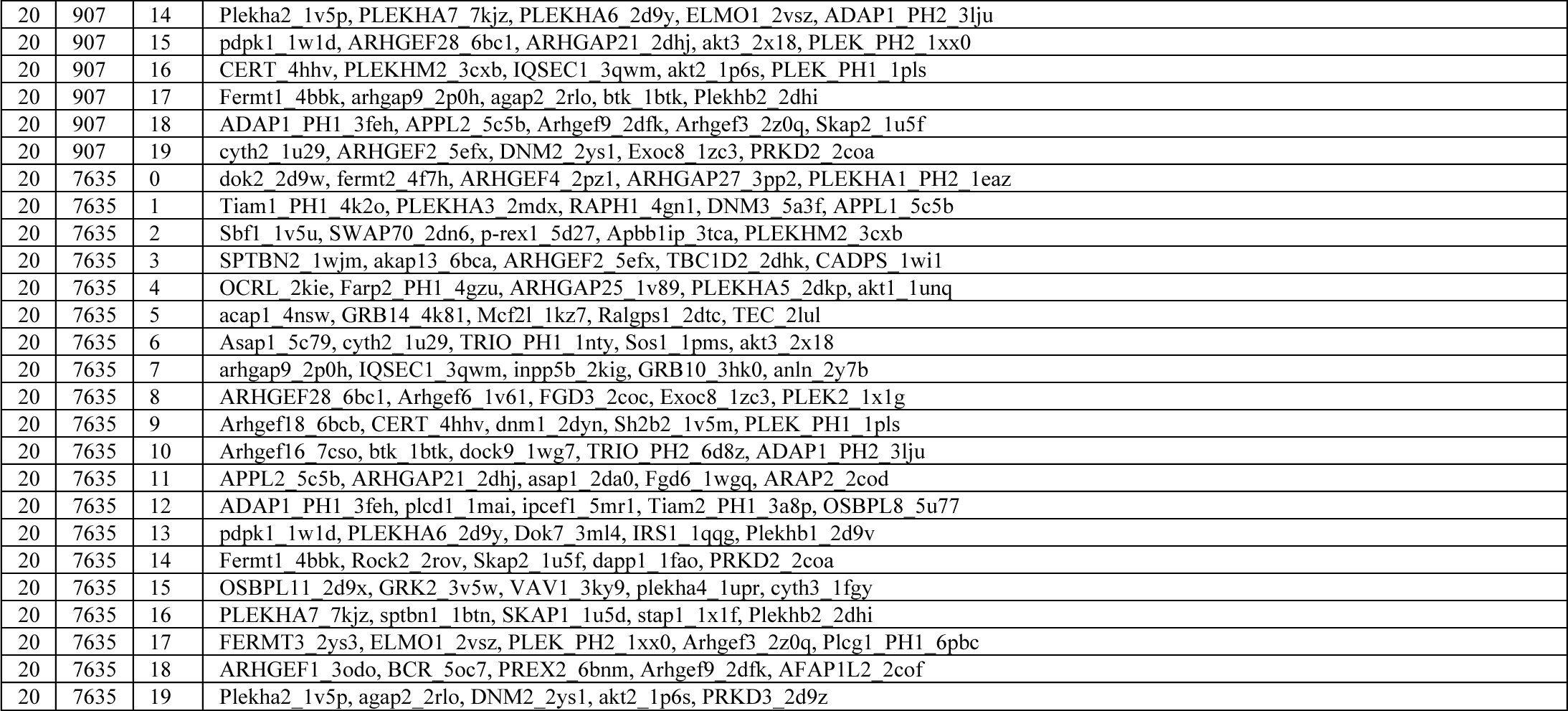
Test set composition for every fold of k-fold cross-validation. Each PH domain is present in only one test set fold for each unique (k, seed) pair.

**Supplementary Table 2.** Test set results for 5-fold cross validation with random seed set to 907.

| 5-fold cross validation, seed = 907 |  |  |  |  |  |  |  |
| --- | --- | --- | --- | --- | --- | --- | --- |
| CV fold number | MSE | Mean Wasserstein Distance | Accuracy (0.8) | Sensitivity (0.8) | Specificity (0.8) | Precision (0.8) | F1 score (0.8) |
| 0 | 0.0221 | 3.86 | 0.954 | 0.71 | 0.97 | 0.6 | 0.65 |
| 1 | 0.0258 | 4.18 | 0.956 | 0.674 | 0.976 | 0.655 | 0.664 |
| 2 | 0.0202 | 2.7 | 0.963 | 0.662 | 0.985 | 0.758 | 0.707 |
| 3 | 0.0253 | 4.64 | 0.965 | 0.696 | 0.983 | 0.733 | 0.714 |
| 4 | 0.0211 | 4.32 | 0.971 | 0.727 | 0.984 | 0.696 | 0.711 |
| **Mean** | **0.0229** | **3.94** | **0.962** | **0.694** | **0.979** | **0.688** | **0.689** |
| **Std** | 0.00253 | 0.749 | 0.00685 | 0.0264 | 0.00645 | 0.0627 | 0.0297 |

**Supplementary Table 3.** Test set results for 5-fold cross validation with random seed set to 7635.

| 5-fold cross validation, seed = 7635 |  |  |  |  |  |  |  |
| --- | --- | --- | --- | --- | --- | --- | --- |
| CV fold number | MSE | Mean Wasserstein Distance | Accuracy (0.8) | Sensitivity (0.8) | Specificity (0.8) | Precision (0.8) | F1 score (0.8) |
| 0 | 0.0225 | 3.64 | 0.961 | 0.771 | 0.975 | 0.708 | 0.738 |
| 1 | 0.021 | 3.74 | 0.957 | 0.738 | 0.972 | 0.641 | 0.686 |
| 2 | 0.0236 | 4.54 | 0.967 | 0.692 | 0.981 | 0.643 | 0.667 |
| 3 | 0.0258 | 3.78 | 0.946 | 0.681 | 0.965 | 0.575 | 0.623 |
| 4 | 0.0254 | 3.85 | 0.957 | 0.806 | 0.966 | 0.578 | 0.673 |
| **Mean** | **0.0237** | **3.91** | **0.957** | **0.738** | **0.972** | **0.629** | **0.677** |
| **Std** | 0.002 | 0.36 | 0.00748 | 0.0527 | 0.00666 | 0.0549 | 0.0411 |

**Supplementary Table 4.** Test set results for 10-fold cross validation with random seed set to 907.

| 10-fold cross validation, seed = 907 |  |  |  |  |  |  |  |
| --- | --- | --- | --- | --- | --- | --- | --- |
| CV fold number | MSE | Mean Wasserstein Distance | Accuracy (0.8) | Sensitivity (0.8) | Specificity (0.8) | Precision (0.8) | F1 score (0.8) |
| 0 | 0.0343 | 6.31 | 0.938 | 0.81 | 0.946 | 0.481 | 0.605 |
| 1 | 0.0259 | 5.49 | 0.955 | 0.716 | 0.97 | 0.615 | 0.662 |
| 2 | 0.0244 | 4.05 | 0.963 | 0.778 | 0.976 | 0.7 | 0.737 |
| 3 | 0.0289 | 4.56 | 0.942 | 0.652 | 0.961 | 0.529 | 0.584 |
| 4 | 0.0218 | 3.18 | 0.958 | 0.69 | 0.978 | 0.705 | 0.696 |
| 5 | 0.021 | 2.85 | 0.96 | 0.789 | 0.972 | 0.667 | 0.723 |
| 6 | 0.0301 | 4.94 | 0.957 | 0.857 | 0.963 | 0.587 | 0.697 |
| 7 | 0.0269 | 3.73 | 0.964 | 0.8 | 0.976 | 0.714 | 0.755 |
| 8 | 0.0214 | 3.64 | 0.964 | 0.78 | 0.974 | 0.622 | 0.692 |
| 9 | 0.0254 | 4.36 | 0.966 | 0.824 | 0.973 | 0.6 | 0.694 |
| **Mean** | **0.026** | **4.31** | **0.957** | **0.77** | **0.969** | **0.622** | **0.684** |
| **Std** | 0.00424 | 1.06 | 0.00963 | 0.0643 | 0.00991 | 0.0772 | 0.0542 |

**Supplementary Table 5.** Test set results for 10-fold cross validation with random seed set to 7635.

| 10-fold cross validation, seed = 7635 |  |  |  |  |  |  |  |
| --- | --- | --- | --- | --- | --- | --- | --- |
| CV fold number | MSE | Mean Wasserstein Distance | Accuracy (0.8) | Sensitivity (0.8) | Specificity (0.8) | Precision (0.8) | F1 score (0.8) |
| 0 | 0.0248 | 3.69 | 0.959 | 0.653 | 0.981 | 0.721 | 0.685 |
| 1 | 0.0183 | 3.81 | 0.961 | 0.793 | 0.974 | 0.714 | 0.751 |
| 2 | 0.0232 | 3.69 | 0.956 | 0.722 | 0.972 | 0.642 | 0.68 |
| 3 | 0.0203 | 3.91 | 0.963 | 0.63 | 0.985 | 0.742 | 0.681 |
| 4 | 0.024 | 4.52 | 0.977 | 0.757 | 0.984 | 0.636 | 0.691 |
| 5 | 0.0243 | 4.94 | 0.958 | 0.731 | 0.973 | 0.636 | 0.681 |
| 6 | 0.0254 | 2.58 | 0.949 | 0.771 | 0.961 | 0.581 | 0.663 |
| 7 | 0.0293 | 5.93 | 0.945 | 0.606 | 0.969 | 0.581 | 0.593 |
| 8 | 0.0246 | 3.96 | 0.964 | 0.785 | 0.975 | 0.662 | 0.718 |
| 9 | 0.0221 | 3.78 | 0.963 | 0.797 | 0.972 | 0.61 | 0.691 |
| **Mean** | **0.0236** | **4.08** | **0.959** | **0.724** | **0.975** | **0.653** | **0.684** |
| **Std** | 0.00297 | 0.89 | 0.00864 | 0.0706 | 0.00731 | 0.057 | 0.0403 |

**Supplementary Table 6.** Test set results for 20-fold cross validation with random seed set to 907.

| 20-fold cross validation, seed = 907 |  |  |  |  |  |  |  |
| --- | --- | --- | --- | --- | --- | --- | --- |
| CV fold number | MSE | Mean Wasserstein Distance | Accuracy (0.8) | Sensitivity (0.8) | Specificity (0.8) | Precision (0.8) | F1 score (0.8) |
| 0 | 0.0318 | 6.17 | 0.949 | 0.765 | 0.962 | 0.578 | 0.658 |
| 1 | 0.0218 | 5.46 | 0.966 | 0.7 | 0.981 | 0.677 | 0.689 |
| 2 | 0.0217 | 2.6 | 0.96 | 0.657 | 0.982 | 0.719 | 0.687 |
| 3 | 0.0229 | 5.63 | 0.958 | 0.69 | 0.975 | 0.629 | 0.657 |
| 4 | 0.0196 | 3.69 | 0.979 | 0.724 | 0.994 | 0.88 | 0.792 |
| 5 | 0.0272 | 3.77 | 0.969 | 0.767 | 0.986 | 0.82 | 0.795 |
| 6 | 0.0352 | 5.09 | 0.955 | 0.667 | 0.974 | 0.629 | 0.647 |
| 7 | 0.0239 | 4.19 | 0.951 | 0.75 | 0.964 | 0.587 | 0.659 |
| 8 | 0.0247 | 2.94 | 0.941 | 0.667 | 0.962 | 0.583 | 0.622 |
| 9 | 0.0233 | 3.3 | 0.98 | 0.816 | 0.992 | 0.886 | 0.849 |
| 10 | 0.0212 | 2.93 | 0.958 | 0.762 | 0.974 | 0.711 | 0.736 |
| 11 | 0.019 | 2.28 | 0.968 | 0.759 | 0.98 | 0.69 | 0.721 |
| 12 | 0.0252 | 4.62 | 0.963 | 0.742 | 0.977 | 0.657 | 0.697 |
| 13 | 0.0253 | 4.72 | 0.957 | 0.81 | 0.966 | 0.591 | 0.684 |
| 14 | 0.0332 | 5.29 | 0.971 | 0.769 | 0.986 | 0.811 | 0.789 |
| 15 | 0.0202 | 2.78 | 0.968 | 0.694 | 0.988 | 0.806 | 0.746 |
| 16 | 0.0212 | 3.62 | 0.969 | 0.852 | 0.975 | 0.639 | 0.73 |
| 17 | 0.0225 | 4.03 | 0.968 | 0.781 | 0.979 | 0.676 | 0.725 |
| 18 | 0.0193 | 3.45 | 0.975 | 0.889 | 0.979 | 0.686 | 0.774 |
| 19 | 0.0202 | 5.23 | 0.974 | 0.75 | 0.985 | 0.692 | 0.72 |
| **Mean** | **0.024** | **4.09** | **0.964** | **0.751** | **0.978** | **0.697** | **0.719** |
| **Std** | 0.00466 | 1.13 | 0.0104 | 0.0619 | 0.00939 | 0.096 | 0.0593 |

**Supplementary Table 7.** Test set results for 20-fold cross validation with random seed set to 7635.

| 20-fold cross validation, seed = 7635 |  |  |  |  |  |  |  |
| --- | --- | --- | --- | --- | --- | --- | --- |
| CV fold number | MSE | Mean Wasserstein Distance | Accuracy (0.8) | Sensitivity (0.8) | Specificity (0.8) | Precision (0.8) | F1 score (0.8) |
| 0 | 0.0244 | 3.86 | 0.96 | 0.756 | 0.978 | 0.756 | 0.756 |
| 1 | 0.0315 | 3.46 | 0.952 | 0.767 | 0.963 | 0.548 | 0.639 |
| 2 | 0.0201 | 3.87 | 0.962 | 0.72 | 0.986 | 0.837 | 0.774 |
| 3 | 0.0184 | 3.7 | 0.972 | 0.81 | 0.982 | 0.743 | 0.776 |
| 4 | 0.0195 | 4.14 | 0.963 | 0.62 | 0.979 | 0.577 | 0.6 |
| 5 | 0.0165 | 3.34 | 0.972 | 0.917 | 0.977 | 0.786 | 0.846 |
| 6 | 0.0206 | 3.42 | 0.958 | 0.744 | 0.974 | 0.69 | 0.716 |
| 7 | 0.028 | 4.26 | 0.956 | 0.794 | 0.966 | 0.574 | 0.667 |
| 8 | 0.0252 | 3.62 | 0.985 | 0.765 | 0.992 | 0.765 | 0.765 |
| 9 | 0.0221 | 4.26 | 0.965 | 0.85 | 0.969 | 0.515 | 0.642 |
| 10 | 0.0314 | 8.31 | 0.961 | 0.778 | 0.973 | 0.651 | 0.709 |
| 11 | 0.0225 | 2.62 | 0.962 | 0.742 | 0.977 | 0.676 | 0.708 |
| 12 | 0.0273 | 4.25 | 0.945 | 0.912 | 0.947 | 0.525 | 0.667 |
| 13 | 0.0287 | 2.47 | 0.94 | 0.722 | 0.956 | 0.553 | 0.627 |
| 14 | 0.0236 | 4.6 | 0.942 | 0.733 | 0.955 | 0.489 | 0.587 |
| 15 | 0.0359 | 6.5 | 0.938 | 0.585 | 0.968 | 0.6 | 0.593 |
| 16 | 0.0219 | 3.54 | 0.967 | 0.808 | 0.975 | 0.618 | 0.7 |
| 17 | 0.0286 | 4.72 | 0.971 | 0.795 | 0.983 | 0.78 | 0.785 |
| 18 | 0.0245 | 3.45 | 0.953 | 0.914 | 0.956 | 0.582 | 0.711 |
| 19 | 0.0249 | 4.13 | 0.967 | 0.75 | 0.976 | 0.581 | 0.655 |
| **Mean** | **0.0248** | **4.13** | **0.96** | **0.774** | **0.972** | **0.642** | **0.696** |
| **Std** | 0.0049 | 1.29 | 0.0119 | 0.0847 | 0.0117 | 0.104 | 0.072 |

**Supplementary Figure S1.**
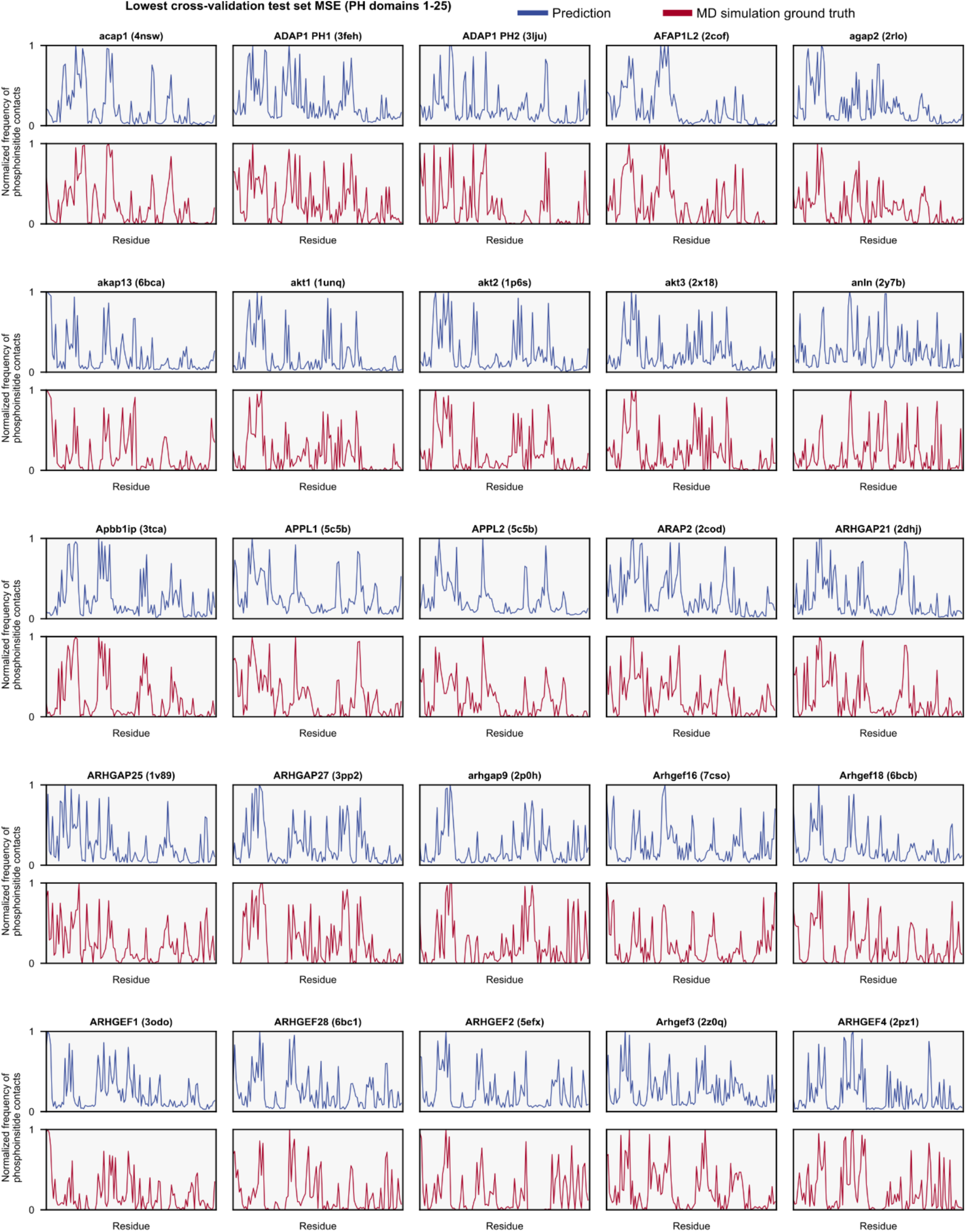
Lowest test set mean squared error predicted normalized phosphoinositide contact frequencies (blue traces) for PH domains 1-25 during cross validation, compared with the results of the reference MD simulations (red traces).

**Supplementary Figure S2.**
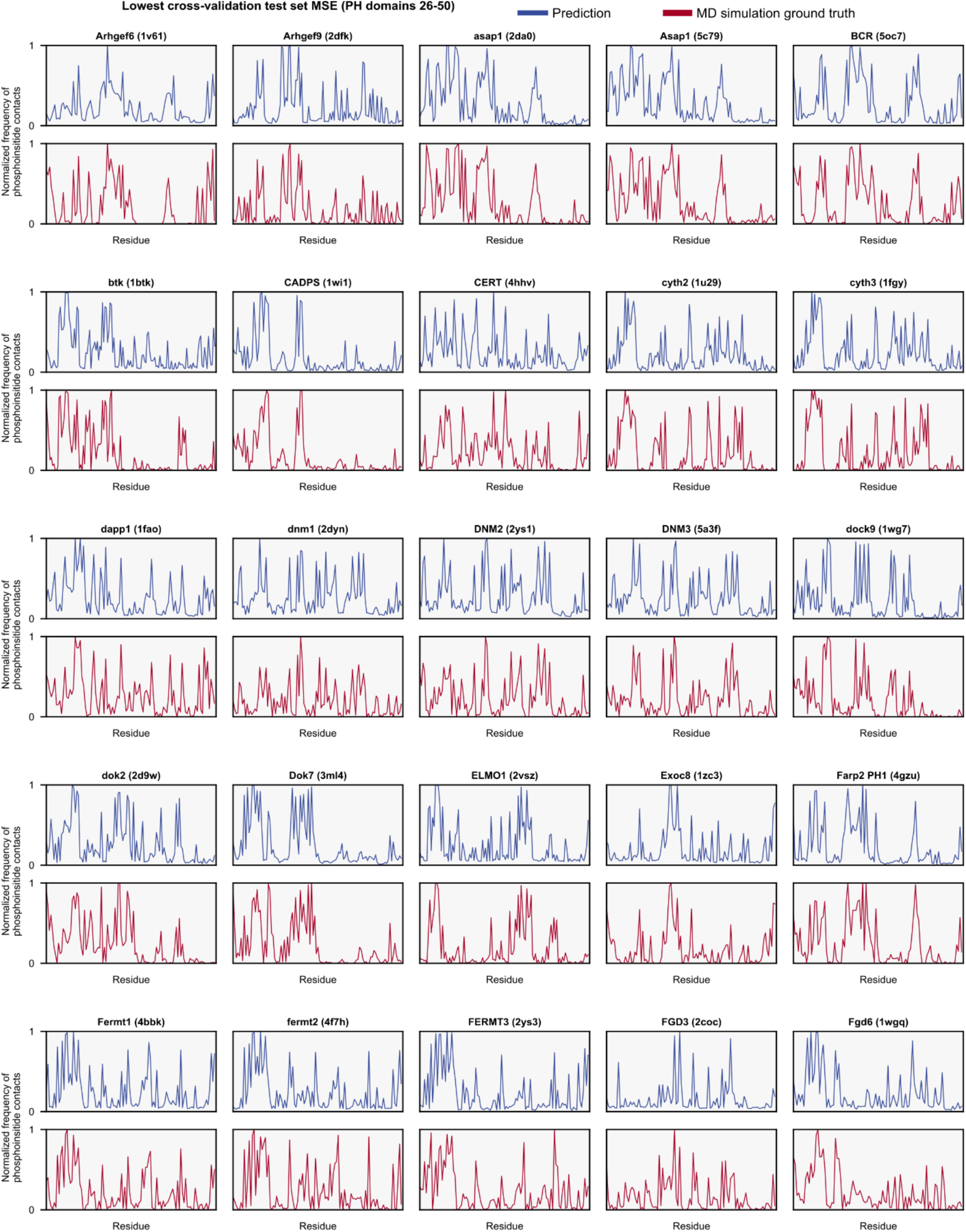
Lowest test set mean squared error predicted normalized phosphoinositide contact frequencies (blue traces) for PH domains 26-50 during cross validation, compared with the results of the reference MD simulations (red traces).

**Supplementary Figure S3.**
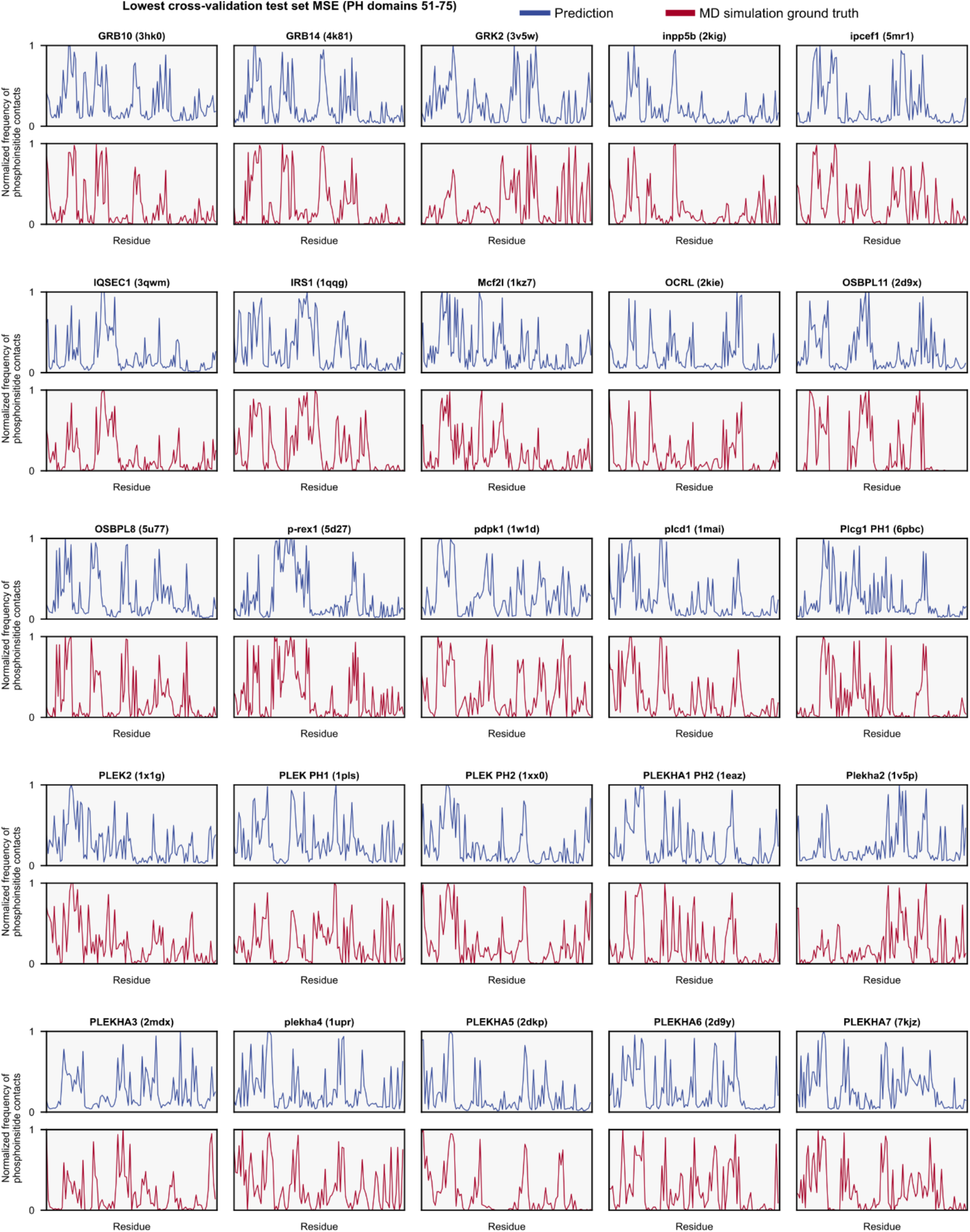
Lowest test set mean squared error predicted normalized phosphoinositide contact frequencies (blue traces) for PH domains 51-75 during cross validation, compared with the results of the reference MD simulations (red traces).

**Supplementary Figure S4.**
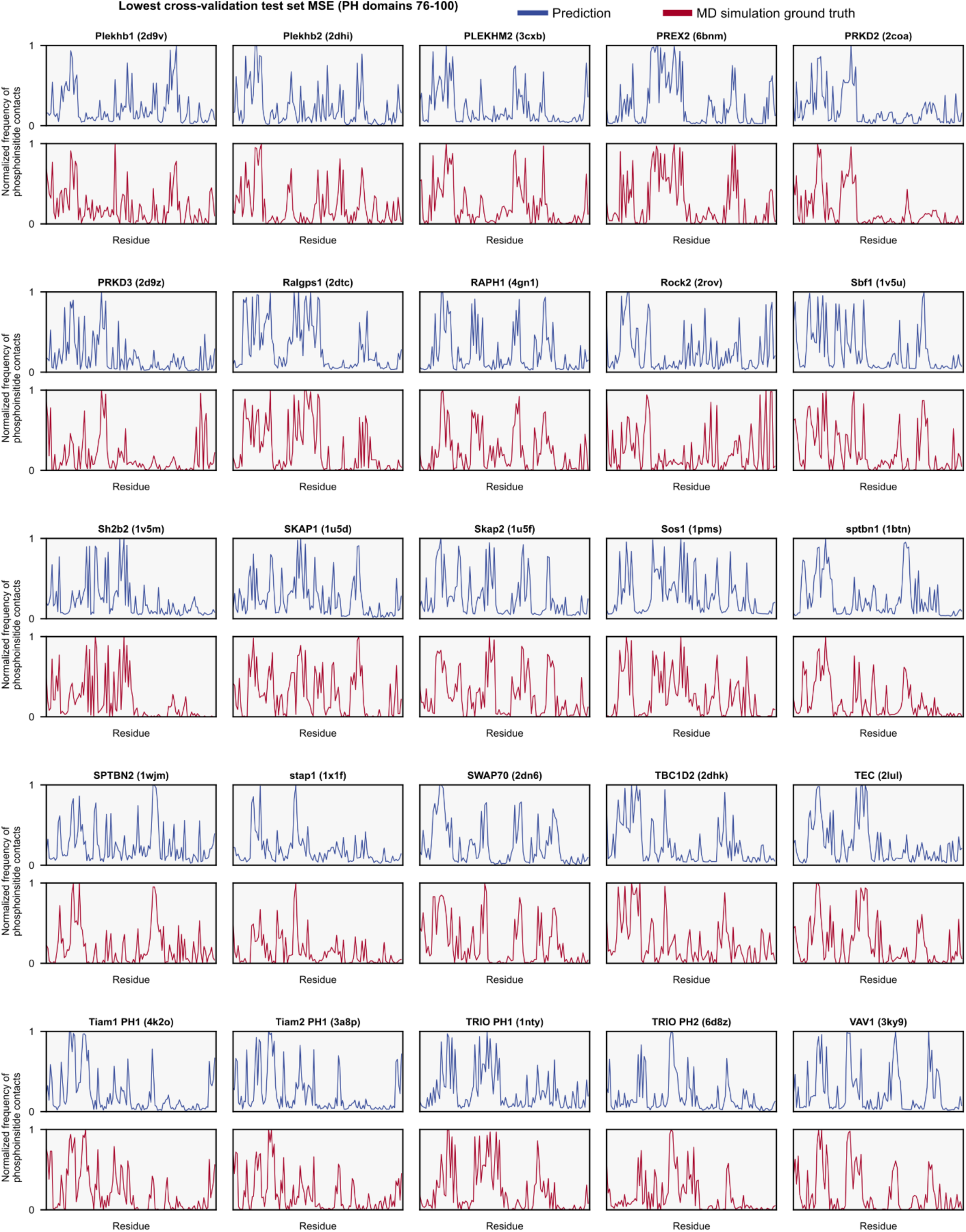
Lowest test set mean squared error predicted normalized phosphoinositide contact frequencies (blue traces) for PH domains 76-100 during cross validation, compared with the results of the reference MD simulations (red traces).

**Supplementary Figure S5.**
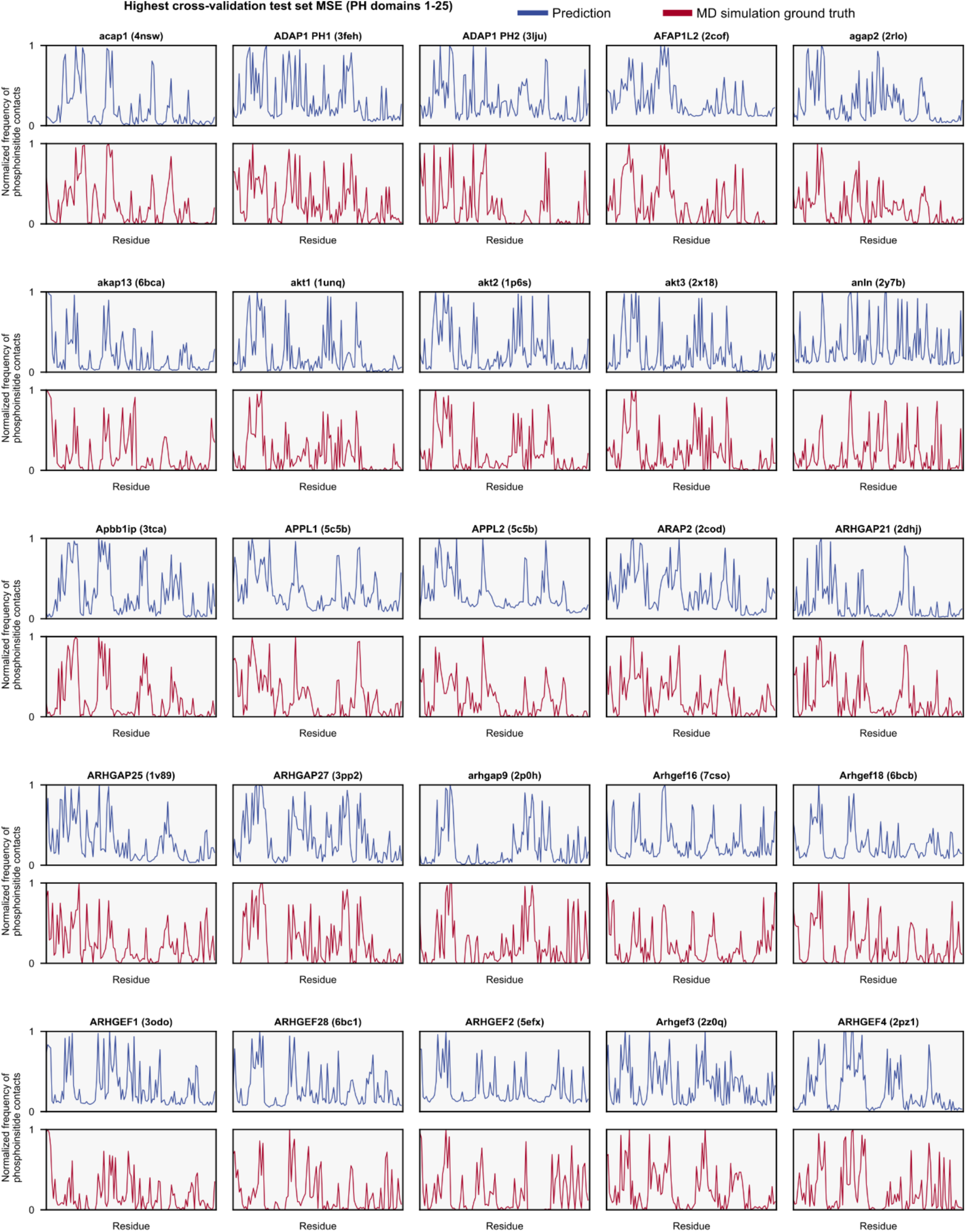
Highest test set mean squared error predicted normalized phosphoinositide contact frequencies (blue traces) for PH domains 1-25 during cross validation, compared with the results of the reference MD simulations (red traces).

**Supplementary Figure S6.**
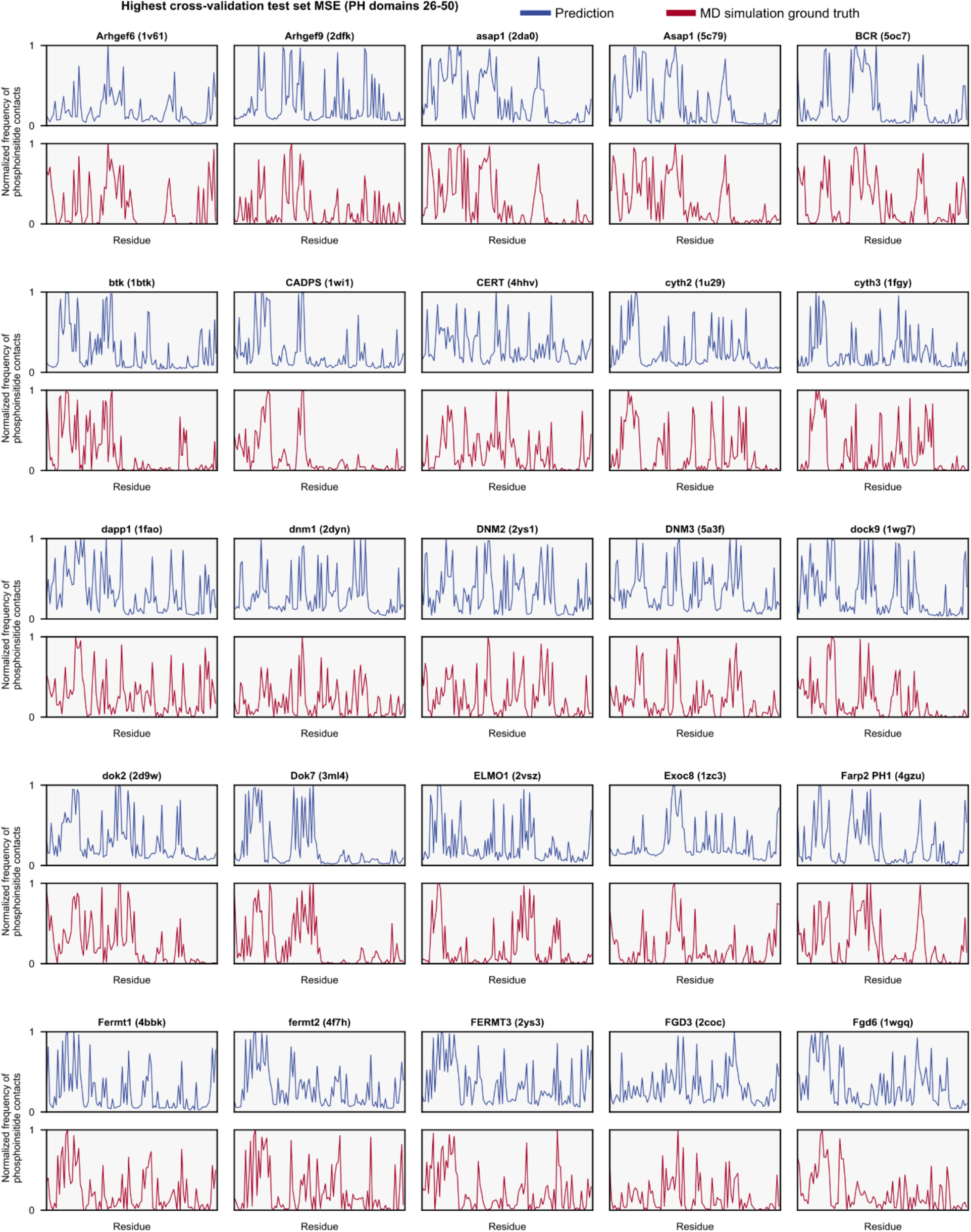
Highest test set mean squared error predicted normalized phosphoinositide contact frequencies (blue traces) for PH domains 26-50 during cross validation, compared with the results of the reference MD simulations (red traces).

**Supplementary Figure S7.**
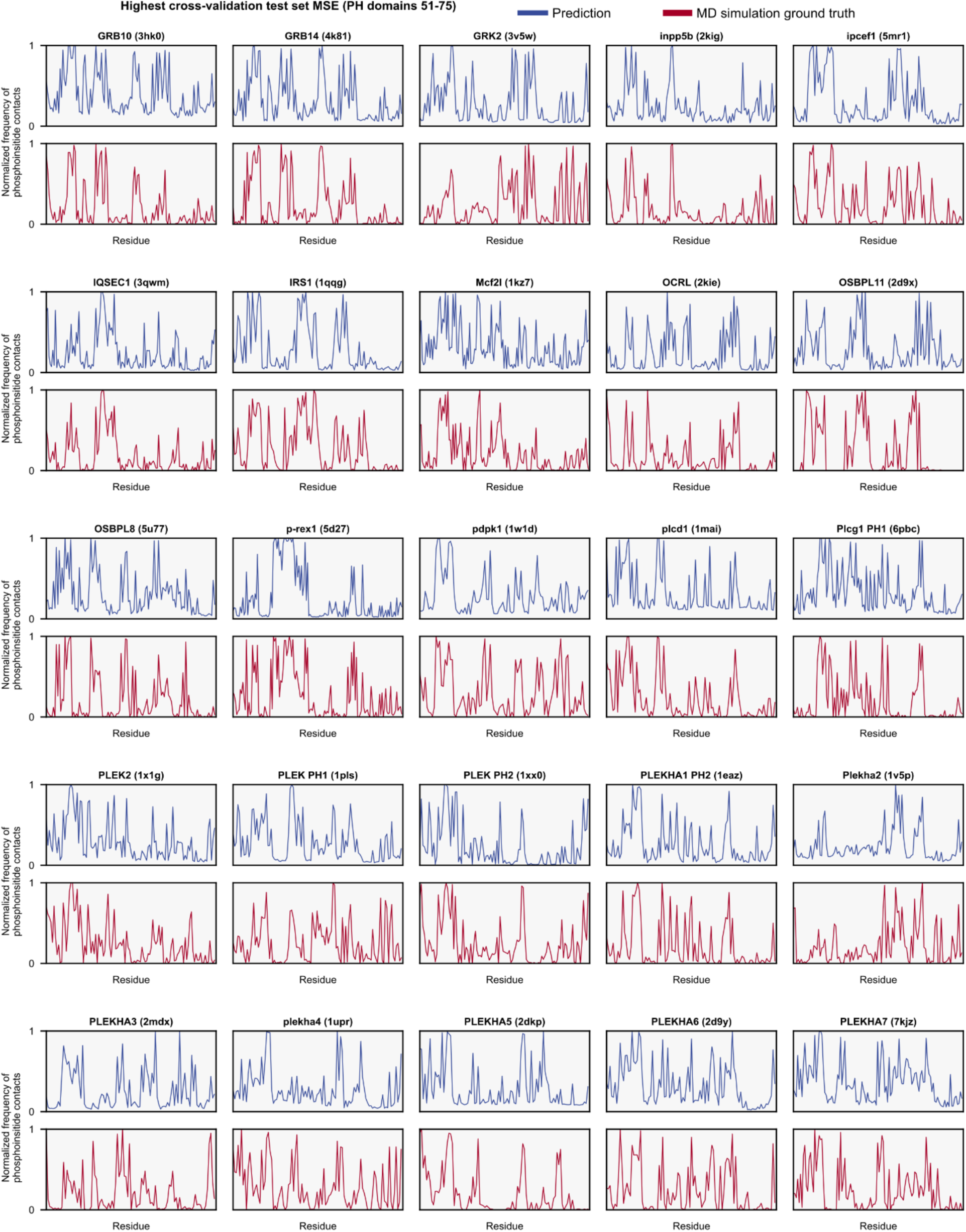
Highest test set mean squared error predicted normalized phosphoinositide contact frequencies (blue traces) for PH domains 51-75 during cross validation, compared with the results of the reference MD simulations (red traces).

**Supplementary Figure S8.**
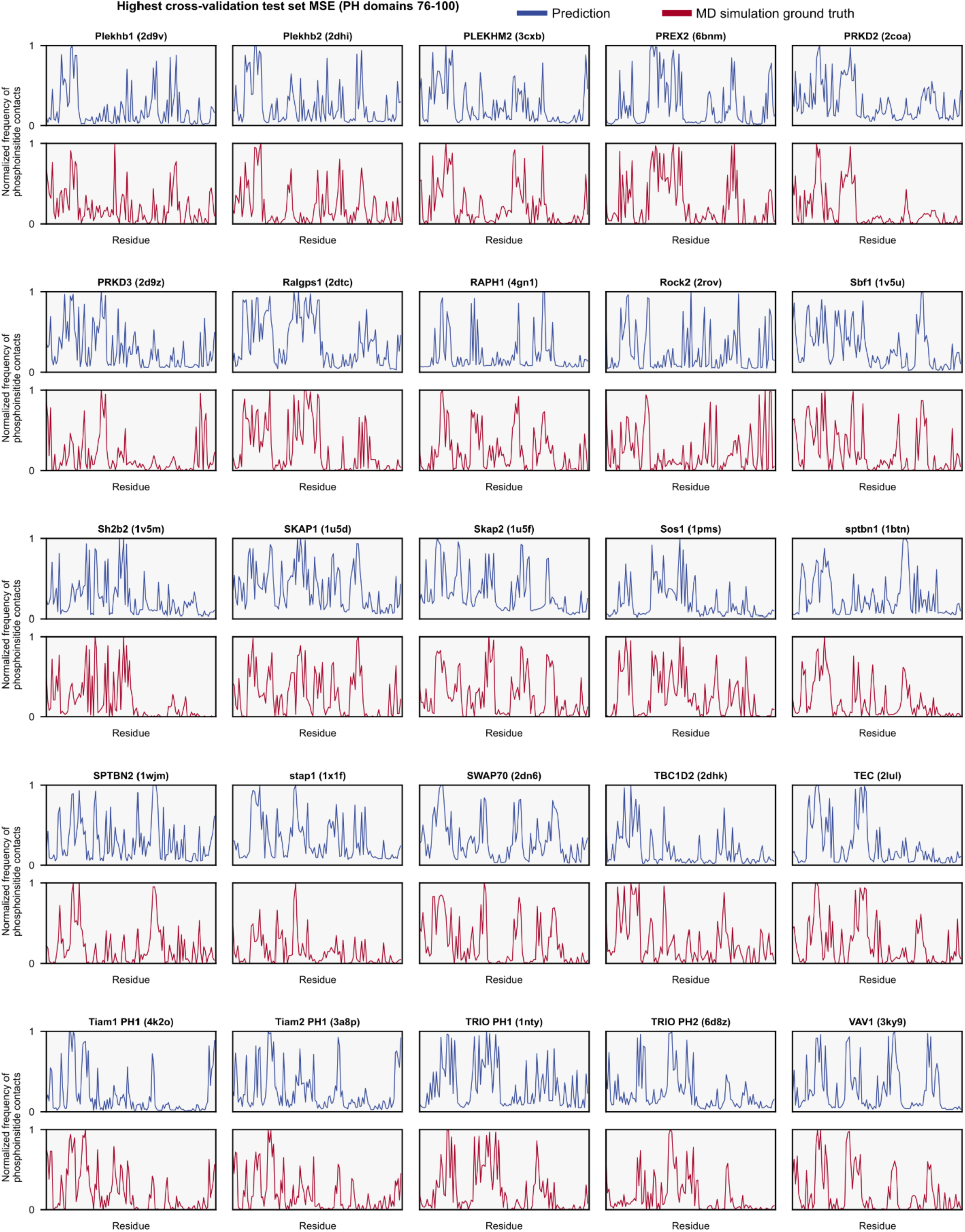
Highest test set mean squared error predicted normalized phosphoinositide contact frequencies (blue traces) for PH domains 76-100 during cross validation, compared with the results of the reference MD simulations (red traces).

**Supplementary Figure S9.**
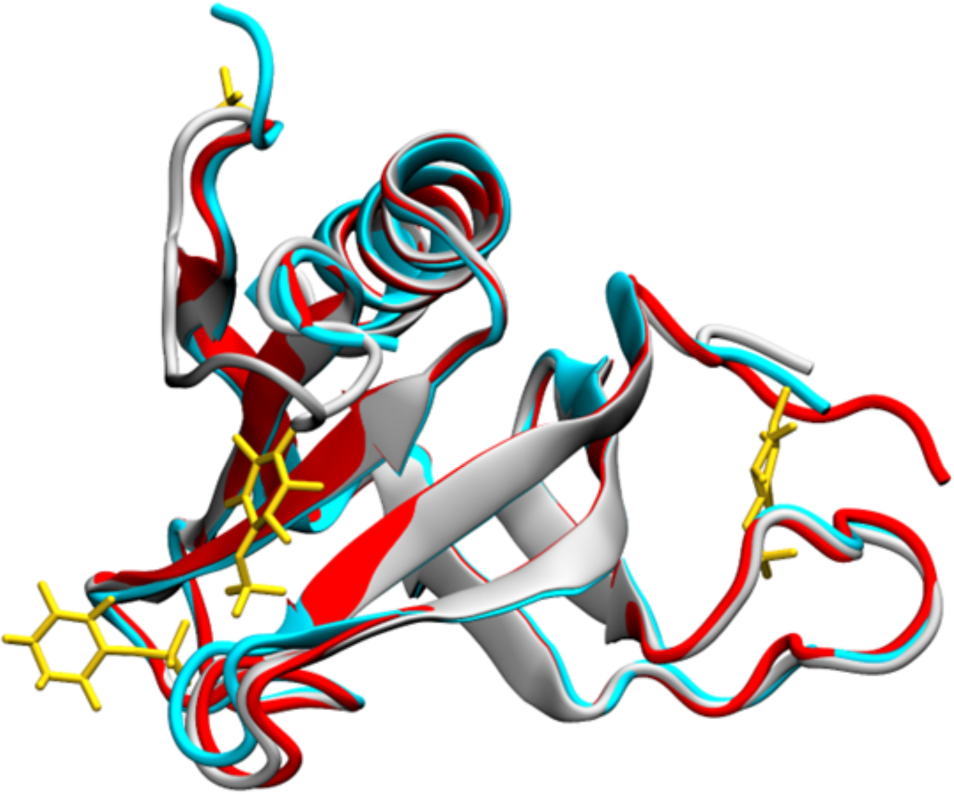
STAMP structural alignment of Yeast Slm1 PH domain structures. PDB ID 4A5K (silver) structure aligned with PDB ID 4A6K (cyan, alignment backbone RMSD = 0.805 Å) and 4A6H (red, alignment backbone RMSD = 0.530 Å). The β5-β6 loop is unresolved in the 4A6K and 4A6H structures, and they display distinct sites of inositol 4-phosphate (yellow sticks) binding.

**Supplementary Figure S10.**
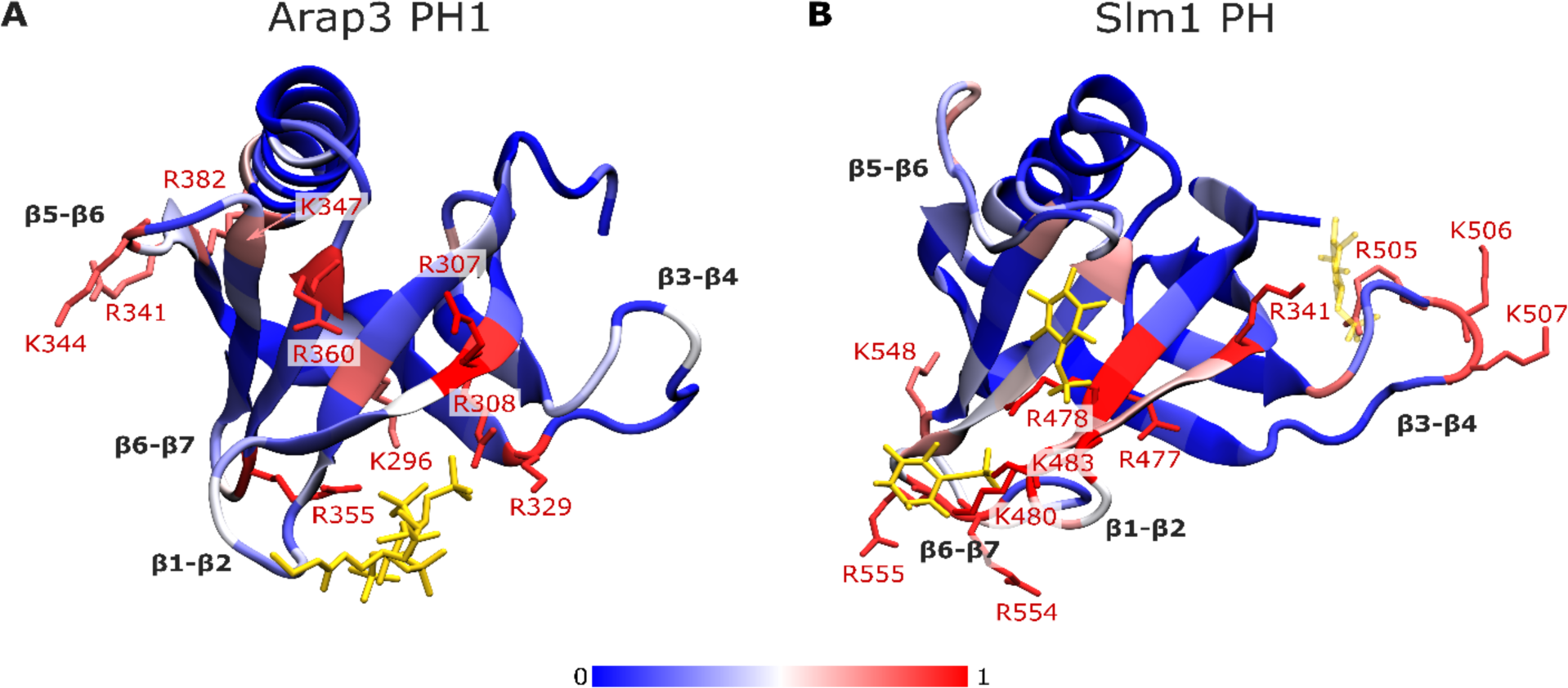
Comparison of predictions with structural data for the Arap3 and Slm1 PH domains. Structures of **A)** human Arap3 PH1 domain (PDB ID: 7YIS) in complex with diC4-PI(3,4,5)P_3_ and **B)** yeast in complex with inositol 4-phosphate. Amino acids are coloured according the predicted normalized frequency of contacts with PIP lipids (see colour scale) predicted by the deep learning model. Residues with predicted normalized frequency of contacts >= 0.8 are displayed as red sticks and labelled. Co-crystallised phosphoinositide analogues and partially resolved phosphate are displayed as yellow sticks. Relevant unstructured loop regions between beta-sheets are labelled. In panel B the unliganded 4A5K protein structure is displayed and used for the prediction, as other structures were missing residues in the β5-β6 loop. STAMP structural alignment was performed between the 4A5K structure and the ligand containing 4A6K (RMSD = 0.805 Å) and 4A6H (RMSD = 0.530 Å) structures, and the inositol 4-phosphate shown are from the these aligned structures. The structural alignments are presented in Supplementary Figure S9.

## Notes

### Competing Interest Statement

The authors have declared no competing interest.

## References

1. A. M. Whited, A. Johs, The interactions of peripheral membrane proteins with biological membranes. Chemistry and Physics of Lipids 192, 51--59 (2015).

2. T. D. Bunney, M. Katan, Phosphoinositide signalling in cancer: beyond PI3K and PTEN. Nature Reviews Cancer 10, 342–352 (2010).

3. M. S. Song, L. Salmena, P. P. Pandolfi, The functions and regulation of the PTEN tumour suppressor. Nature Reviews Molecular Cell Biology 13, 283–296 (2012).

4. B. Vanhaesebroeck, L. Stephens, P. Hawkins, PI3K signalling: the path to discovery and understanding. Nature Reviews Molecular Cell Biology 13, 195–203 (2012).

5. L. H. Wong, A. T. Gatta, T. P. Levine, Lipid transfer proteins: the lipid commute via shuttles, bridges and tubes. Nature Reviews Molecular Cell Biology 20, 85–101 (2019).

6. J. Dudek, Role of cardiolipin in mitochondrial signaling pathways. Frontiers in Cell and Developmental Biology 5, 1--17 (2017).

7. V. Teixeira, M. J. Feio, M. Bastos, Role of lipids in the interaction of antimicrobial peptides with membranes. Progress in Lipid Research 51, 149–177 (2012).

8. S. Bleicken et al., Structural Model of Active Bax at the Membrane. Molecular Cell 56, 496--505 (2014).

9. A. Arkhipov, Y. Yin, K. Schulten, Four-Scale Description of Membrane Sculpting by BAR Domains. Biophysical Journal 95, 2806–2821 (2008).

10. J. C. Porta et al., Molecular architecture of the human caveolin-1 complex. Science Advances 8, eabn7232 (2022).

11. Andreas H. Larsen, Laura H. John, Mark S. P. Sansom, Robin A. Corey, Specific interactions of peripheral membrane proteins with lipids: what can molecular simulations show us? Bioscience Reports 42, BSR20211406 (2022).

12. W. Cho, R. V. Stahelin, Membrane-Protein Interactions in Cell Signaling and Membrane Trafficking. Annual Review of Biophysics and Biomolecular Structure 34, 119–151 (2005).

13. B. Antonny, Mechanisms of Membrane Curvature Sensing. Annual Review of Biochemistry 80, 101–123 (2011).

14. C. Grauffel et al., Cation−π Interactions As Lipid-Specific Anchors for Phosphatidylinositol-Specific Phospholipase C. Journal of the American Chemical Society 135, 5740–5750 (2013).

15. S. Pant, E. Tajkhorshid, Microscopic Characterization of GRP1 PH Domain Interaction with Anionic Membranes. Journal of Computational Chemistry 41, 489--499 (2020).

16. K. E. Landgraf, C. Pilling, J. J. Falke, Molecular mechanism of an oncogenic mutation that alters membrane targeting: Glu17Lys modifies the PIP lipid specificity of the AKT1 PH domain. Biochemistry 47, 12260--12269 (2008).

17. K. M. Ferguson, M. A. Lemmon, J. Schlessinger, P. B. Sigler, Structure of the high affinity complex of inositol trisphosphate with a phospholipase C pleckstrin homology domain. Cell 83, 1037–1046 (1995).

18. T. Harayama, H. Riezman, Understanding the diversity of membrane lipid composition. Nature Reviews Molecular Cell Biology 19, 281--296 (2018).

19. C. G. Sudhahar, R. M. Haney, Y. Xue, R. V. Stahelin, Cellular membranes and lipid-binding domains as attractive targets for drug development. Curr Drug Targets 9, 603–613 (2008).

20. D. M. Boes, A. Godoy-Hernandez, D. G. G. McMillan, Peripheral Membrane Proteins: Promising Therapeutic Targets across Domains of Life. Membranes. 2021 (10.3390/membranes11050346).

21. K. Segers et al., Design of protein–membrane interaction inhibitors by virtual ligand screening, proof of concept with the C2 domain of factor V. Proceedings of the National Academy of Sciences 104, 12697–12702 (2007).

22. Z. Liu et al., Trp2313-His2315 of factor VIII C2 domain is involved in membrane binding: structure of a complex between the C2 domain and an inhibitor of membrane binding. J Biol Chem 285, 8824–8829 (2010).

23. B. Miao et al., Small molecule inhibition of phosphatidylinositol-3,4,5-triphosphate (PIP3) binding to pleckstrin homology domains. Proc Natl Acad Sci U S A 107, 20126–20131 (2010).

24. A. Nawrotek et al., PH-domain-binding inhibitors of nucleotide exchange factor BRAG2 disrupt Arf GTPase signaling. Nature Chemical Biology 15, 358–366 (2019).

25. J. L. Scott, C. A. Musselman, E. Adu-Gyamfi, T. G. Kutateladze, R. V. Stahelin, Emerging methodologies to investigate lipid-protein interactions. Integr Biol (Camb) 4, 247–258 (2012).

26. V. Corradi et al., Emerging Diversity in Lipid-Protein Interactions. Chemical Reviews 119, 5775–5848 (2019).

27. A. W. Smith, Lipid-protein interactions in biological membranes: A dynamic perspective. Biochimica et Biophysica Acta - Biomembranes 1818, 172--177 (2012).

28. G. van Meer, A. I. de Kroon, Lipid map of the mammalian cell. J Cell Sci 124, 5–8 (2011).

29. G. van Meer, D. R. Voelker, G. W. Feigenson, Membrane lipids: where they are and how they behave. Nat Rev Mol Cell Biol 9, 112–124 (2008).

30. V. Biou, Lipid-membrane protein interaction visualised by cryo-EM: A review. Biochimica et Biophysica Acta (BBA) - Biomembranes 1865, 184068 (2023).

31. J. F. van Dyck, A. Konijnenberg, F. Sobott, in Membrane Protein Structure and Function Characterization: Methods and Protocols, J.-J. Lacapere, Ed. (Springer New York, New York, NY, 2017), pp. 205–232.

32. K. Gupta et al., Identifying key membrane protein lipid interactions using mass spectrometry. Nature protocols 13, 1106--1120 (2018).

33. H. Y. Yen et al., PtdIns(4,5)P2 stabilizes active states of GPCRs and enhances selectivity of G-protein coupling. Nature 559, 423--427 (2018).

34. J. E. Keener, G. Zhang, M. T. Marty, Native Mass Spectrometry of Membrane Proteins. Analytical Chemistry 93, 583–597 (2021).

35. C. Sahin, D. J. Reid, M. T. Marty, M. Landreh, Scratching the surface: native mass spectrometry of peripheral membrane protein complexes. Biochemical Society Transactions 0, 1--12 (2020).

36. M. Frick, C. Schwieger, C. Schmidt, Liposomes as Carriers of Membrane-Associated Proteins and Peptides for Mass Spectrometric Analysis. Angewandte Chemie - International Edition 60, 11523--11530 (2021).

37. G. Hedger, M. S. P. Sansom, Lipid interaction sites on channels, transporters and receptors: Recent insights from molecular dynamics simulations. Biochimica et Biophysica Acta - Biomembranes 1858, 2390--2400 (2016).

38. F. Naughton, Interactions of peripheral membrane proteins with phosphatidylinositol lipids: insights from molecular dynamics simulations. (2017).

39. M. P. Muller et al., Characterization of Lipid-Protein Interactions and Lipid-Mediated Modulation of Membrane Protein Function through Molecular Simulation. Chemical Reviews 119, 6086--6161 (2019).

40. S. J. Marrink et al., Computational Modeling of Realistic Cell Membranes. Chemical Reviews 119, 6184--6226 (2019).

41. A. Buyan et al., Piezo1 Forms Specific, Functionally Important Interactions with Phosphoinositides and Cholesterol. Biophysical Journal 119, 1683–1697 (2020).

42. F. X. Contreras et al., Molecular recognition of a single sphingolipid species by a protein’s transmembrane domain. Nature 481, 525--529 (2012).

43. M. Manna, Niemel, Mechanism of allosteric regulation of $\beta$2-adrenergic receptor by cholesterol. eLife 5, 1--21 (2016).

44. M. R. Schmidt, P. J. Stansfeld, S. J. Tucker, M. S. P. Sansom, Simulation-based prediction of phosphatidylinositol 4,5-bisphosphate binding to an ion channel. Biochemistry 52, 279--281 (2013).

45. E. Pyle et al., Structural Lipids Enable the Formation of Functional Oligomers of the Eukaryotic Purine Symporter UapA. Cell Chemical Biology 25, 840–848.e844 (2018).

46. K. Zhou et al., A Ceramide-Regulated Element in the Late Endosomal Protein LAPTM4B Controls Amino Acid Transporter Interaction. ACS Central Science 4, 548–558 (2018).

47. O. Soubias et al., Membrane surface recognition by the ASAP1 PH domain and consequences for interactions with the small GTPase Arf1. Science Advances 6, (2020).

48. S. Dadsena et al., Ceramides bind VDAC2 to trigger mitochondrial apoptosis. Nature Communications 10, 1832 (2019).

49. K. I. P. Le Huray, T. D. Bunney, N. Pinotsis, A. C. Kalli, M. Katan, Characterization of the membrane interactions of phospholipase Cγ reveals key features of the active enzyme. Science Advances 8, eabp9688 (2022).

50. K. I. P. Le Huray, H. Wang, F. Sobott, A. C. Kalli, Systematic simulation of the interactions of pleckstrin homology domains with membranes. Science Advances 8, eabn6992 (2022).

51. E. Yamamoto et al., Multiple lipid binding sites determine the affinity of PH domains for phosphoinositide-containing membranes. Science Advances 6, (2020).

52. G. Hedger, M. S. P. Sansom, Kolds, The juxtamembrane regions of human receptor tyrosine kinases exhibit conserved interaction sites with anionic lipids. Scientific Reports 5, 9198 (2015).

53. E. Yamamoto, A. C. Kalli, K. Yasuoka, M. S. P. Sansom, Interactions of Pleckstrin Homology Domains with Membranes: Adding Back the Bilayer via High-Throughput Molecular Dynamics. Structure 24, 1421--1431 (2016).

54. T. D. Newport, M. S. P. Sansom, P. J. Stansfeld, The MemProtMD database: A resource for membrane-embedded protein structures and their lipid interactions. Nucleic Acids Research 47, D390--D397 (2019).

55. R. A. Corey et al., Identification and assessment of cardiolipin interactions with E. coli inner membrane proteins. Science Advances 7, eabh2217 (2021).

56. M. A. Lemmon, Membrane recognition by phospholipid-binding domains. Nature Reviews Molecular Cell Biology 9, 99--111 (2008).

57. M. Lenoir, I. Kufareva, R. Abagyan, M. Overduin, Membrane and Protein Interactions of the Pleckstrin Homology Domain Superfamily. Membranes 5, 646--663 (2015).

58. N. Singh et al., Redefining the specificity of phosphoinositide-binding by human PH domain-containing proteins. Nature Communications 12, 4339 (2021).

59. P. Garcia et al., The Pleckstrin Homology Domain of Phospholipase C-$\delta$1 Binds with High Affinity to Phosphatidylinositol 4,5-Bisphosphate in Bilayer Membranes. Biochemistry 34, 16228--16234 (1995).

60. K. Anand, K. Maeda, A. C. Gavin, Structural analyses of the Slm1-PH domain demonstrate ligand binding in the non-canonical site. PLoS ONE 7, 1--10 (2012).

61. D. F. J. Ceccarelli et al., Non-canonical interaction of phosphoinositides with pleckstrin homology domains of Tiam1 and ArhGAP9. Journal of Biological Chemistry 282, 13864--13874 (2007).

62. Y. Liu, R. A. Kahn, J. H. Prestegard, Interaction of fapp1 with Arf1 and PI4P at a membrane surface: An example of coincidence detection. Structure 22, 421--430 (2014).

63. X. Jian et al., Molecular Basis for Cooperative Binding of Anionic Phospholipids to the PH Domain of the Arf GAP ASAP1. Structure 23, 1977--1988 (2015).

64. R. A. Kahn, D. G. Lambright. (2015).

65. A. E. Aleshin et al., Structural basis for the association of PLEKHA7 with membrane-embedded phosphatidylinositol lipids. bioRxiv, 1--16 (2020).

66. S. B. T. A. Amos, A. C. Kalli, J. Shi, M. S. P. Sansom, Membrane Recognition and Binding by the Phosphatidylinositol Phosphate Kinase PIP5K1A: A Multiscale Simulation Study. Structure 27, 1336--1346.e1332 (2019).

67. M. Chavent et al., Interactions of the EphA2 Kinase Domain with PIPs in Membranes: Implications for Receptor Function. Structure 26, 1025–1034.e1022 (2018).

68. P. Veličković et al., Graph Attention Networks. 2017 (10.48550/arXiv.1710.10903).

69. S. Brody, U. Alon, E. Yahav, How Attentive are Graph Attention Networks? 2021 (10.48550/arXiv.2105.14491).

70. A. Vaswani et al., Attention is all you need. Advances in neural information processing systems 30, (2017).

71. J. Devlin, M.-W. Chang, K. Lee, K. Toutanova, Bert: Pre-training of deep bidirectional transformers for language understanding. arXiv preprint arXiv:1810.04805, (2018).

72. OpenAI, GPT-4 Technical Report. 2023 (10.48550/arXiv.2303.08774).

73. A. Dosovitskiy, et al., An image is worth 16×16 words: Transformers for image recognition at scale. *arXiv preprint arXiv:*2010.11929, (2020).

74. R. D. King, O. I. Orhobor, C. C. Taylor, Cross-validation is safe to use. Nature Machine Intelligence 3, 276–276 (2021).

75. L. Wang, J. Zhang, D. Wang, C. Song, Membrane contact probability: An essential and predictive character for the structural and functional studies of membrane proteins. PLOS Computational Biology 18, e1009972 (2022).

76. Y. Zhang et al., Structural Insights Uncover the Specific Phosphoinositide Recognition by the PH1 Domain of Arap3. International Journal of Molecular Sciences 24, 1125 (2023).

77. O. Gallego et al., A systematic screen for proteing-lipid interactions in Saccharomyces cerevisiae. Molecular Systems Biology 6, (2010).

78. J. Jumper et al., Highly accurate protein structure prediction with AlphaFold. Nature 596, 583–589 (2021).

79. A. Kume et al., The Pleckstrin Homology Domain of Diacylglycerol Kinase &#x3b7; Strongly and Selectively Binds to Phosphatidylinositol 4,5-Bisphosphate *. Journal of Biological Chemistry 291, 8150–8161 (2016).

80. S. Schurmans, S. Polizzi, A. Scoumanne, S. Sayyed, P. Molina-Ortiz, The Ras/Rap GTPase activating protein RASA3: From gene structure to in vivo functions. Advances in Biological Regulation 57, 153–161 (2015).

81. P. J. Lockyer et al., Distinct subcellular localisations of the putative inositol 1,3,4,5-tetrakisphosphate receptors GAP1IP4BP and GAP1m result from the GAP1IP4BP PH domain directing plasma membrane targeting. Current Biology 7, 1007–1010 (1997).

82. G. E. Cozier et al., GAP1 IP4BP Contains a Novel Group I Pleckstrin Homology Domain That Directs Constitutive Plasma Membrane Association. Journal of Biological Chemistry 275, 28261–28268 (2000).

83. H. Li et al., Phosphoinositide Conversion Inactivates R-RAS and Drives Metastases in Breast Cancer. Advanced Science 9, 2103249 (2022).

84. E. R. Smith, L. R. Caulley, A. R. Storm, A. M. Hulse-Kemp, A. K. Stoeckman, Gossypium hirsutum gene of unknown function Gohir.A03G007700.1 encodes a potential VAN3-binding protein with a phosphoinositide-binding site. MicroPubl Biol 2023, (2023).

85. M. Chandra et al., Classification of the human phox homology (PX) domains based on their phosphoinositide binding specificities. Nature Communications 10, (2019).

86. C.-Z. Zhou et al., Crystal Structure of the Yeast Phox Homology (PX) Domain Protein Grd19p Complexed to Phosphatidylinositol-3-phosphate *. Journal of Biological Chemistry 278, 50371–50376 (2003).

87. Z. Jia, R. Ghai, B. M. Collins, A. E. Mark, The recognition of membrane-bound PtdIns3P by PX domains. Proteins: Structure, Function, and Bioinformatics 82, 2332–2342 (2014).

88. A. Chatzigoulas, Z. Cournia, Predicting protein–membrane interfaces of peripheral membrane proteins using ensemble machine learning. Briefings in Bioinformatics 23, bbab518 (2022).

89. N. van Hilten, J. Methorst, N. Verwei, H. J. Risselada, Physics-based generative model of curvature sensing peptides; distinguishing sensors from binders. Science Advances 9, eade8839 (2023).

90. F. C. Chollet, et al. (2015).

91. Robert T. McGibbon et al., MDTraj: A Modern Open Library for the Analysis of Molecular Dynamics Trajectories. Biophysical Journal 109, 1528–1532 (2015).

92. N. Michaud-Agrawal, E. J. Denning, T. B. Woolf, O. Beckstein, MDAnalysis: A toolkit for the analysis of molecular dynamics simulations. Journal of Computational Chemistry 32, 2319–2327 (2011).

93. R. Gowers et al., paper presented at the Proceedings of the Python in Science Conference, 2016.

94. D. P. Kingma, J. Ba, Adam: A Method for Stochastic Optimization. 2014 (10.48550/arXiv.1412.6980).

95. P. Virtanen et al., SciPy 1.0: fundamental algorithms for scientific computing in Python. Nature Methods 17, 261–272 (2020).

